# Finding the right balance of conservation effort in cultivated grasslands: A modelling study on protecting dispersers in a climatically changing and anthropogenically disturbed environment

**DOI:** 10.1101/2022.09.20.508676

**Authors:** Johannes A. Leins

## Abstract

Managing cultivated grasslands in a sustainable way is controversial, because it often goes along with economical loss and additional effort for local farmers. On the plus side, such a management could permit inhabiting species not only to survive but to thrive and expand their range. In order to satisfy both aspects, it can be helpful to minimize conservation effort to a degree that is still ecologically beneficial but intervenes as little as possible with regional land-use customs. Computer simulations are a useful tool to find such compromises prior to implementing management strategies. We simulated the population development of the large marsh grasshopper, a grassland species with limited dispersal abilities, in a disturbed and climatically changing environment of Germany up to the year 2080. Our results show that - in a spatially aggregated landscape - adapting the harvesting schedule in a relatively low number ≤ 7 % of (in)directly connected yet otherwise intensively managed grasslands suffices for species preservation and even expansion to some extent. The effect on dispersal success of additional conservation effort above this 7 % threshold is significantly lower than it is below the threshold. In terms of population size, however, every additional refuge benefits the grasshopper. Climate change enhances the positive effects on the target species even further. A higher level of fragmentation, however, requires a substantially larger conservation effort in terms of protected grassland proportion. Therefore, it is recommended and more effective to focus on the implementation of protected areas within spatially aggregated grasslands. Stakeholders should additionally be aware of the fact that it can take several years for a conservation effort to become apparent and measurable, especially if the goal is to support an isolated or reintroduced species in expanding into unpopulated territories.

## 1 Introduction

Unsuitable land use practices can amplify the negative impact of global warming or constrain the adaptive capacities of endemic species (Oliver & Morecroft, 2014). In fact, there are instances of formerly endangered species benefiting from climate change in theory that could still be prevented from thriving by regional land use practices (Leins et al., 2021; Leins et al., 2022; Poniatowski et al., 2018a). From a perspective of conservation planning, it is imperative to identify measures that support target species or ecological communities on a long run before implementing them, especially in cultivated landscapes.

In such regions, it is impossible to implement comprehensive measures at will to achieve a conservation goal, as land is often in private ownership or other interests are of (higher) relevance. Rather, it is necessary to take focused and metered conservation measures that allow target species to thrive and, ideally, expand their range despite the disturbed environment. With the respective knowledge it can be easier to either find the land owners’ acceptance towards conservation measures or intervene with their land use practices as little as possible (Moloney et al., 2018; Will et al., 2021; Nguyen et al., 2022). As thoroughly discussed in our previous studies (Leins et al., 2021; Leins et al., 2022), simulation models together with population viability analysis (PVA) are a valuable tool to aid stakeholders in their effort of identifying such suitable measures. We showed furthermore that suitably managing smaller grassland plots can suffice to support populations locally and that even less suitable, homogeneously distributed management plans can allow moderate dispersal of a species.

However, both natural and cultivated environments are usually more heterogeneous in terms of composition and usage. Projecting required connectivity between suitable habitats to allow successful dispersal in an otherwise disturbed environment is more challenging. There are different concepts on assessing connectivity in randomly distributed environments. In percolation theory (Stauffer & Aharony, 1994), a critical probability threshold is determined above which the general connectivity of an (infinite) environment is assured. On basis of this theory, With (2002) suggested a proportion threshold level ≥ 50 % of connected replicate landscapes (generated using the same probability value) as a more reasonable measure for applied movement ecology to assess likely connectivity in finite landscapes. Another approach extends the binary definition of suitable and unsuitable habitats to include habitats of intermediate suitability (Wiegand et al., 2005; Wiegand et al., 1999). These so-called poor-quality habitats could function as stepping stones between suitable habitats that are otherwise (too) far apart, and in this way achieve connectivity way below the thresholds mentioned before.

All above concepts are considered for the analysis and setup of the present study using another extension of the HiLEG model (Supplement S1). The study explores the effects of applying conservation effort of increasing extent (number of protected habitats) on the population development and dispersal success of the large marsh grasshopper (LMG, *Stethophyma grossum)* in cultivated grasslands of different fragmentation levels. More precisely, the analysis aims at identifying the effort required to support an established LMG population at the edge of uninhabited territory in dispersing into new habitats of North Germany depending on projected climate change scenarios (CCS) of increasing severity. We addressed this issue with the following research questions:

1. How does the relative effort in conservation-oriented grassland management affect the population development of a species with limited dispersal ability?
2. Are there time-critical factors that are worth considering for conservation planning in a climatically changing environment?
3. Does the conservation effort required to meet a conservation target differ depending on the spatial landscape structure?

The formerly endangered (Winkler & Haacks, 2019; Winkler, 2000) LMG is a species well suited for such an analysis. It is native to wet grasslands (Heydenreich, 1999) and is affected differently by external factors such as climatic conditions (Wingerden et al., 1991; Ingrisch, 1983) and land use (Leins et al., 2021) during its annual life cycle. Studies confirm that in theory it is benefiting from global warming (Leins et al., 2021; Poniatowski et al., 2018a; Trautner & Hermann, 2008), but at the same time its range could mostly remain restricted by land use practices (Löffler et al., 2019; Poniatowski et al., 2018a; Leins et al., 2022). It occasionally traverses greater distances, but its basic dispersal ability is rather low rendering it vulnerable to local disturbances. Mowing schedules that could allow reasonable regional development of the species exist (Leins et al., 2021; Marzelli, 1997), yet, the broad implementation of such schedules could prove difficult due to their reduced cost-effectiveness (Gerling et al., 2022).

Particularly regarding the latter difficulty, the present study intends to evaluate a limiting configuration of regional land use schedules. That is, a simulation setup in which single protected refuges are randomly distributed (with varying probability) in an intensively managed environment, i.e., a limited number of refuges in cultivated grasslands of high yield. Local populations must therefore cope in an environment that is suitable in principle, but for the most part highly disturbed. Taking into account two spatial configurations of a landscape, the grasslands surrounding an initially isolated population is either aggregated with a high number of (suitable) habitats, or fragmented with a low number of respective habitats. Both the spatial configuration and the location of the initial populations are obtained from realistic data and surveys of North German grasslands. As another factor affecting the LMG’s life cycle, three CCS of increasing severity were applied during simulation runs.

Overall, the simulation results are expected to clarify, if and to what extent configurations of minimal heterogeneity (high number of intensively managed grassland versus varying, yet low number of refuges) already suffice to achieve a regionally sustained or expanding LMG population.

## 2 Material and Methods

The experimental setup (Section 2.1) and evaluation parameters (Section 2.2) used for the present study are described in the following. Table 1 gives and overview of the relevant parameters for initialization and evaluation of the simulation runs. The simulations were run using the most recent version of the HiLEG model^1^ and the output data^2^,^3^ as well as calculated evaluation data^4^ are available online. For a detailed description of the implementation and parameterization of the HiLEG model, please refer to Supplement.

**Table 1:** Selection of parameters used to initialize a simulation run (above double line) and to evaluate simulation results (below). First column: parameter name used in text. Second column: mathematical parameter symbol. Third column: valid parameter value(s) and their units (if applicable). Fourth column: brief description of parameter.

| Parameter Name | Symbol | Value(s) / Unit | Description |
| --- | --- | --- | --- |
| starting date | $t_{init}$ | 01 January 2020 | The date of initial time steps translated to climate data index |
| duration | $t_{\Delta}$ | 21,840 *days* | Runtime in days resp. time steps |
| habitat area | $A_{hab}$ | $250 \times 250 \text{ m}^2$ | Area of a grassland habitat |
| initial region | $reg_{init}$ | $\in \{aggr, frag\}$ | Definition of originating habitat (region in terms of spatial configuration of grassland surrounding it, where *aggr*=aggregated and *frag*=fragmented) |
| climate change scenario | $CCS$ | $\in \{FF, MOD, BAU\}$ | Representative Concentration Pathways of $CO_2$ model |
| start of vegetation period | $t_{veg}$ | day | Day on which the yearly sum of surface temperature $\omega_{ts}$ reaches $200.0^\circ\text{C}$ (see Eqn. 1) |
| mowing schedule | $T_{mow}$ | $\in \{T_{conv}, T_{prot}\}$ | Applied set of yearly mowing events at a distinct grassland habitat |
| conventional mowing schedule | $T_{conv}$ | $= \{42, 84, 126, 168, 210\}$ | Yearly timing (days after $t_{veg}$) of mowing events in conventional managed grasslands |
| protective mowing schedule | $T_{prot}$ | $= \{49, 217\}$ | Yearly timing (days after $t_{veg}$) of mowing events in protected habitats |
| protected grassland probability | $p_{prot}$ | $\in \{0.01, 0.02, \dots, 0.2, 1.0\}$ | Probability of grassland to be defined as protected habitat ($T_{mow} = T_{prot}$) at simulation start |
| dispersal radius | $rad_{disp}$ | 1,500 *m* | Maximum distance covered by an individual (Griffioen, 1996) |
| inhabited status | $stat_{inh}$ | $\in \{occ, est, res\}$ | The inhabited status of a grassland habitat, where *occ*=occupied, *est*=established, *res*=residential |
| potential range | $rng_{pot}$ | *m* | The distance in meters from habitat of origin to farthest habitat (in)directly connected by $rad_{disp}$ |
| realized (inhabited) range | $rng_{inh},$<br>$inh \in stat_{inh}$ | *m* | The distance in meters from habitat of origin to farthest occupied / established / residential habitat |
| number of changing habitats | $\Delta n_{inh},$<br>$inh \in stat_{inh}$ | $\in \mathbb{N}$ | Yearly number of habitats changing their inhabited status to occupied / established / residential |
| population density | $dens_{occ}$ | *ind. m*<sup>2</sup> | Population density in *individuals m*<sup>2</sup> considering all occupied habitats in the region |

Abbreviations: *aggr*=aggregated, *BAU*=business as usual, *CCS*=climate change scenario, *conv*=conventional, *dens*=density, *disp*=dispersal, *est*=established, *FF*=full force, *frag*=fragmented, *hab*=habitat, *inh*=inhabited, *init*=initial, *m*=meters, *MOD*=moderate, *mow*=mowing, *occ*=occupied, *pot*=potential, *prot*=protective / protected, *rad*=radius, *reg*=region, *res*=residential, *rng*=range, *scen*=scenario, *stat*=status, *t*=time step, *veg*=vegetation

### 2.1 Experimental Setup

We used the stage- and cohort-based, spatially explicit HiLEG model to simulate the development of an LMG population dispersing from one out of two known grassland habitats on the edge of grasslands in the German federal state of Schleswig-Holstein (SH) (Fig. 2A-B) that are currently uninhabited by the LMG. Following Griffioen (1996) the LMG’s maximum dispersal radius (potentially connecting two habitats) was defined as *rad_disp_* = 1,500 *m* and the occasional long distance dispersal (Oppel, 2005) due to its principle flight ability (Sörens, 1996) was ignored for the present analysis. Existing grasslands were subdivided into habitat plots with an area of *A_hab_* = 250 × 250 *m*^2^ each. The first known habitat (in the upper center of the state) is located in a landscape of spatially rather aggregated grasslands, while the surroundings of the second known habitat (on the northern border of the state) are highly fragmented, yet still within dispersal range of the target species. For reasons of comparability, only one of these habitats is initially considered populated in a single simulation run. This originating habitat is defined as protected, i.e., biotic and abiotic conditions (despite climate) are considered ideal, and only low-impact grassland mowing at the start and end of the vegetation period (cf. Table 1, *T_prot_*) is applied to account for management required to maintain a favorable vegetation structure for the LMG (Marzelli, 1997). Apart from that, mowing has a solely lethal impact on the population, but to a significantly different extent depending on the species’ life stage (Leins et al., 2021). Surrounding grasslands are either exposed to a conventional mowing schedule with five cuts per year (cf. Table 1, *T_conv_*), or randomly selected to function as protected habitat similar to the habitats of origin. The probability *p_prot_* to be selected as protected habitat is defined at simulation start, where possible values are *p_prot_* ∈ {0.01, 0.02,…, 0.2, 1.0}. Here, a simulation with *p_prot_* = 1.0 functions as control or benchmark and represents a scenario where all habitats are defined as protected.

The timing of grassland mowing is coupled to the start of the vegetation period *t_veg_*. Following Gerling et al. (2020), this period starts when the yearly temperature sum surpasses 200.0 °*C*. Adapting their calculation to the surface temperature *ω_ts_* used in the present study gives the following equation:

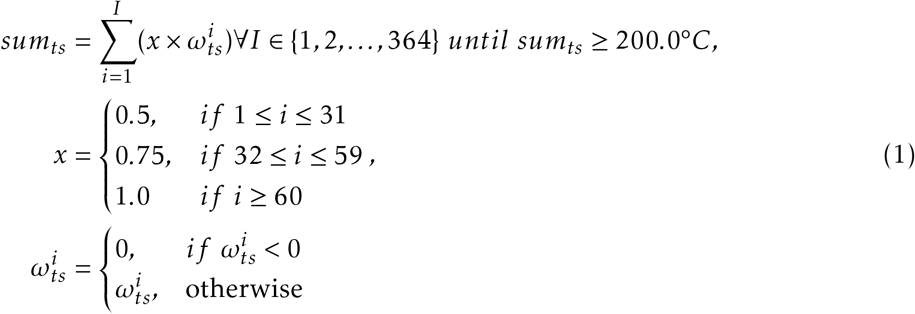

Here, *sum_ts_* is the summed surface temperature, 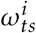 is the mean surface temperature (ignoring negative values) on day *i* of a year, and *x* is a weight including the temperature values of January and February with only 50 % and 75 % of their extent. The value of *t_veg_* equals the day *i* where *t_sum_* reaches 200.0 °*C*.

Simulations run for 60 years on the basis of a daily time step starting January 2020 and ending December 2079. One out of three CCS is applied at simulation start. Below, these scenarios will be distinguished by action taken towards reducing *CO*_2_ emissions: full force (FF, RCP2.6), moderate (MOD, RCP4.5) and business as usual (BAU, RCP8.5), where RCP stands for *Representative Concentration Pathways of CO*_2_. The projected climate parameters of daily resolution have a different effect on the processes of population dynamics depending on the current life stage (e.g. Wingerden et al., 1991; Ingrisch, 1983) and define the start of the vegetation period as described above. They also differ on the spatial scale and at least slightly per habitat.

For each combination of the above simulation parameters, 100 replicates were created, with each of them using a unique random seed. Per replicate, or rather random seed, this leads to a different distribution of protected habitats, time series of climate projections and changes the outcome of stochastic processes on a cohort-level.

### 2.2 Evaluation Parameters

To determine dispersal success and population development depending on simulation setup, we calculated or extracted several evaluation parameters from the simulation output and grouped them by the initialization parameters (Table 1) *reg_init_* ∈ {*aggr, frag*} (initial region containing the habitat of origin), *T_mow_* ∈ {*T_conv_, T_prot_*} (conventional / protective mowing schedule), *CCS* ∈ {*FF,MOD,BAU*} (climate change scenario) and *p_prot_* ∈ {0.01, 0.02,…, 0.2, 1.0} (protected grassland probability). The results were further accumulated by simulation year and spatially distinguished by two states of a grassland habitat: (1) the mowing schedule randomly determined at simulation start (*T_mow_* ∈ {*T_conv_,T_prot_*}), where conventional usage (*T_conv_*) represents a five-cut mowing schedule and protective usage (*T_prot_*) defines a species’ *refuge* with two cuts at start and end of the vegetation period (cf. Table 1 for definition of mowing days); and (2) the inhabited status during a simulation year (*stat_inh_* ∈ {*occ, est, res*}), i.e., whether a habitat was *occ*upied at all, at some point contained a large enough population to be considered established when monitored (*imago density* ≥ 0.002 *individuals m*^-2^, including immigrants), or at some point contained a large enough residential population (hatched from eggs laid in preceding year, i.e., *imago density* ≥ 0.002 *individuals m*^-2^, excluding immigrants). Figure 1 illustrates the inhabited status of a hypothetical grassland cell using a stylized representation of imago density development within three years. The distinction between the inhabited status is crucial for the interpretation of the results: a habitat may become randomly occupied for a brief period, but never develop a substantial population size; locally measuring a large enough (thus theoretically established) population can be due to a high number of immigrants from nearby (protected) habitats; considering a population residential highlights a locally uninterrupted life cycle, but as a measure it masks the presence of smaller populations.

**Figure 1:**
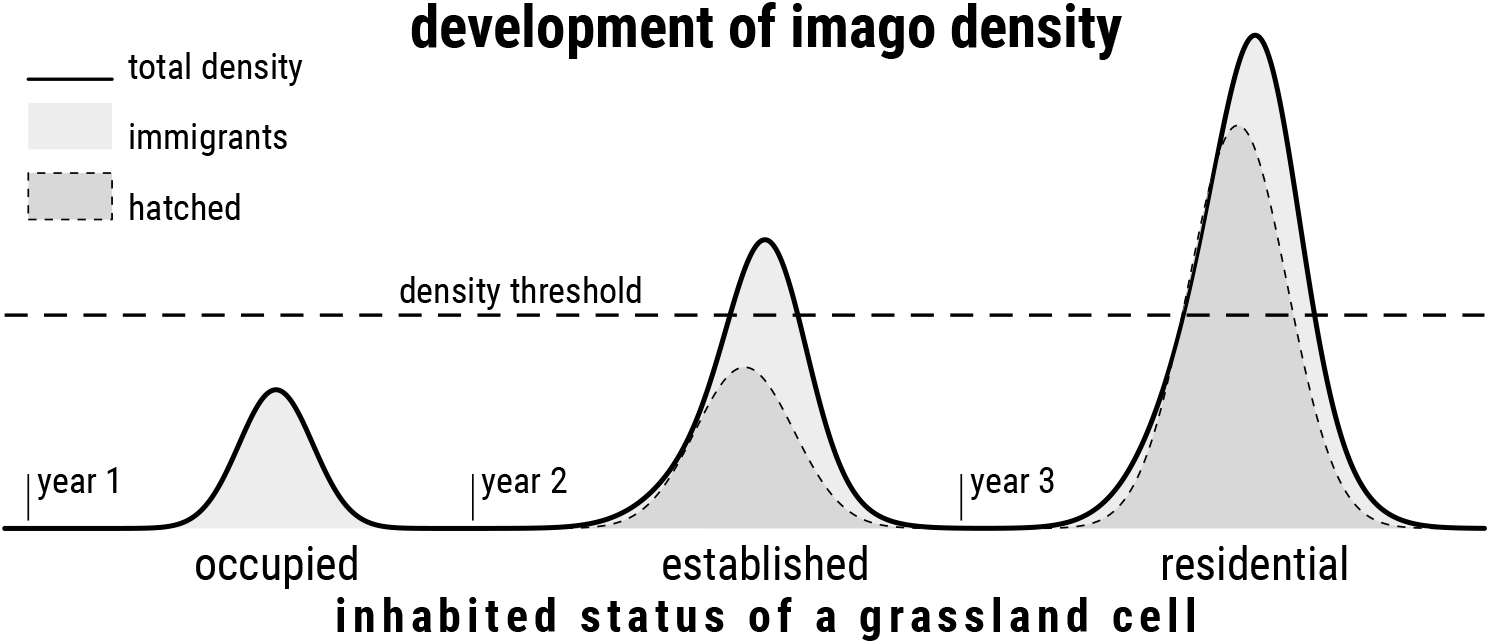
Stylized development of imago density in a hypothetical grassland cell within three years to illustrate the local inhabited status. The solid line represents the current total imago density, the light grey areas the density share of immigrants, and the dark grey areas, framed by a dashed line, the density share originating from eggs hatched in the cell itself. The dashed horizontal line marks the density threshold responsible for a change in inhabited status: if a total density > 0 remains below the threshold, the cell is considered *occupied*; if total density exceeds the threshold, the status changes to *established*; if, in addition, the density share of hatched eggs exceeds the threshold, the local population is considered *residential*

The evaluation parameters used in the analysis have two levels. One is the spatial configuration that arises from the random distribution of protected habitats. Here, two protected habitats are defined as directly connected if the distance between them is not greater than the LMG’s dispersal radius (*rad_disp_* = 1,500 *m*). If there is a connection through other directly connected protected habitats, the two habitats in question are considered to be indirectly connected. All protected habitats (in)directly connected to the habitat of origin are considered the *protected network*. The straight distance [meters] from the originating habitat to the farthest habitat in the protected network will be called *potential range* (*range_pot_*). When ignoring the habitats’ land use type (*type_use_*), all grasslands (in)directly connected to the originating habitat are considered the *functional network* and described as *functionally connected*.

The second level of evaluation parameters is the realized distribution and development of an LMG population depending on the categories and spatial configuration described above: (1) the straight distance [meters] from the habitat containing the initial population to the farthest occupied habitat (*realized occupied range, range_occ_*), established habitat (*realized established range, range_est_*) or residential habitat (*realized residential range, range_res_*); (2) the yearly number of habitats changing their inhabited status to occupied (Δ*n_occ_*), established (Δ*n_est_*) or residential (Δ*n_res_*) for the first time; and (3) the mean population density [*individuals m*^2^] of all occupied habitats (*dens_occ_*).

## 3 Results

Benchmark for the analysis are the replicate(s) that yielded the most optimistic results by the end of the simulation in 2079. These were the ones parameterized with ideal (yet unrealistic) conditions of 100 % protected grasslands (*p_prot_* = 1.0) in the most severe scenario BAU, as the LMG benefits from global warming (Leins et al., 2021). Here, the grasshopper managed to occupy habitats in distances of up to 13,313 *m* (aggregated region) and 8,139 *m* (fragmented region). The resulting distribution under ideal conditions is depicted in Figure 2B (black dots).

**Figure 2:**
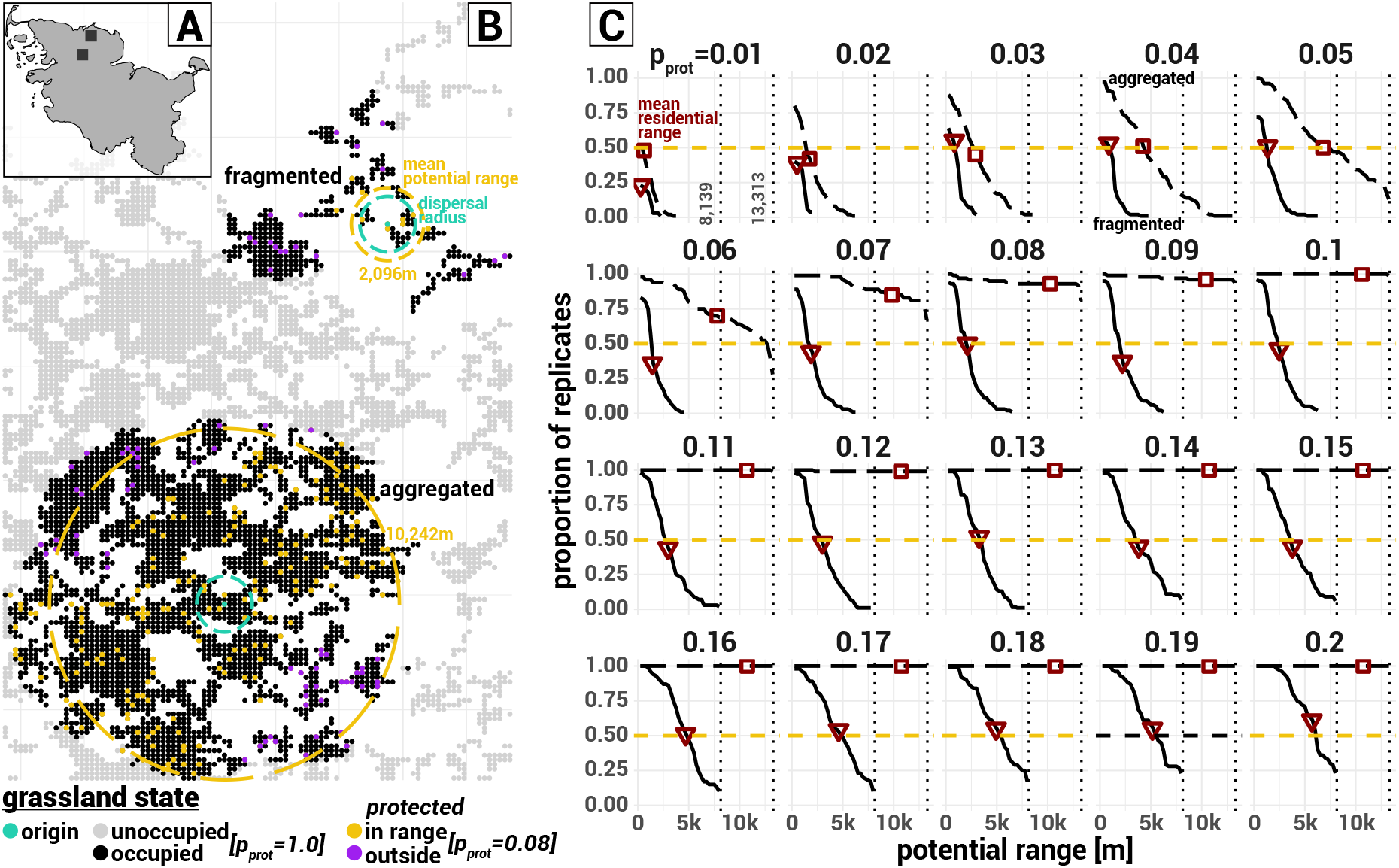
Outline map of the federal state Schleswig-Holstein (A), with the black rectangles marking both study regions containing the habitats of origin. Distribution of grasslands (grey dots) in the study region (B) and habitats occupied at simulation end (black dots) under ideal conditions (*p_prot_* = 1.0, RCP8.5) in at least 1 out of 100 replicates. Eeach dot represents a grassland plot with an area of 62,500 *m*^2^. The green dots mark the habitats of origin in the fragmented and aggregated landscape, and the green circle their dispersal radius. Other colored dots highlight the distribution of protected grasslands for one realization of *p_prot_* = 0.08, where the orange dots mark refuges belonging to the protected network of either of the originating habitats, and the purple dots the ones outside the network. The orange circles depict the potential ranges of the same *p_prot_* value depending on region (fragmented: 2,096 *m*; aggregated: 10,242 *m*). Proportion of replicates (y-axis) having a respective minimum potential range (x-axis) from the originating habitat’s perspective (C) in the fragmented (solid black lines) or aggregated region (dashed black lines) depending on the protected grassland probability *p_prot_* ranging from 0.01 to 0.2 (subplots). The horizontal orange dashed line marks a proportion of 0.5. Red marks highlight the mean realized residential range at simulation end depending on region (triangle: fragmented; square: aggregated)

The potential range (cf. Section 2.2) is different depending on initial region. Figure 2B (colored dots) shows the distribution of protected habitats exemplary for one realization (or replicate) of *P_prot_* = 0.08, where the orange dots represent habitats belonging to the *protected network* (orange dots) in either one of the initial regions. In general, the proportion of replicates having protected habitats within a certain *potential range* greatly differs between regions (Figure 2C). While for the aggregated grasslands and *p_prot_* > 0.06 in more than 50 % of the cases (cf. *threshold level* in With, 2002) the potential range reaches as far as the realized occupied range under ideal conditions (Figure 2C, black dashed lines above purple horizontal line), this is only true for a small percentage ≪ 50 % of replicates for *p_prot_* > 0.13 of the fragmented landscape (Figure 2C, black solid lines).

Contrary, the proportion (or number) of replicates having certain realized (residential) ranges (Figure 3, Table 2A-C) does not match the potential range. The LMG’s residential range is usually below or occasionally equal to the determined potential range of protected grasslands (Figures 3C and F, Table 2A). In none of the simulations (protected) grasslands outside the protected network received sufficient immigration to allow an established or residential population (Figures 3B-C and E-F, purple line). This is despite the observation that there are a occasions during the simulation runs where grasshoppers occupy grasslands outside the reach of the closest refuge (Figures 3A and D, dots above purple line; Table 2C), but in numbers too low to survive without repeated additional immigration. Thus, the functional connectivity of grasslands (independent of their inhabited status) did not result in realized (residential) ranges outside the protected network.

**Figure 3:**
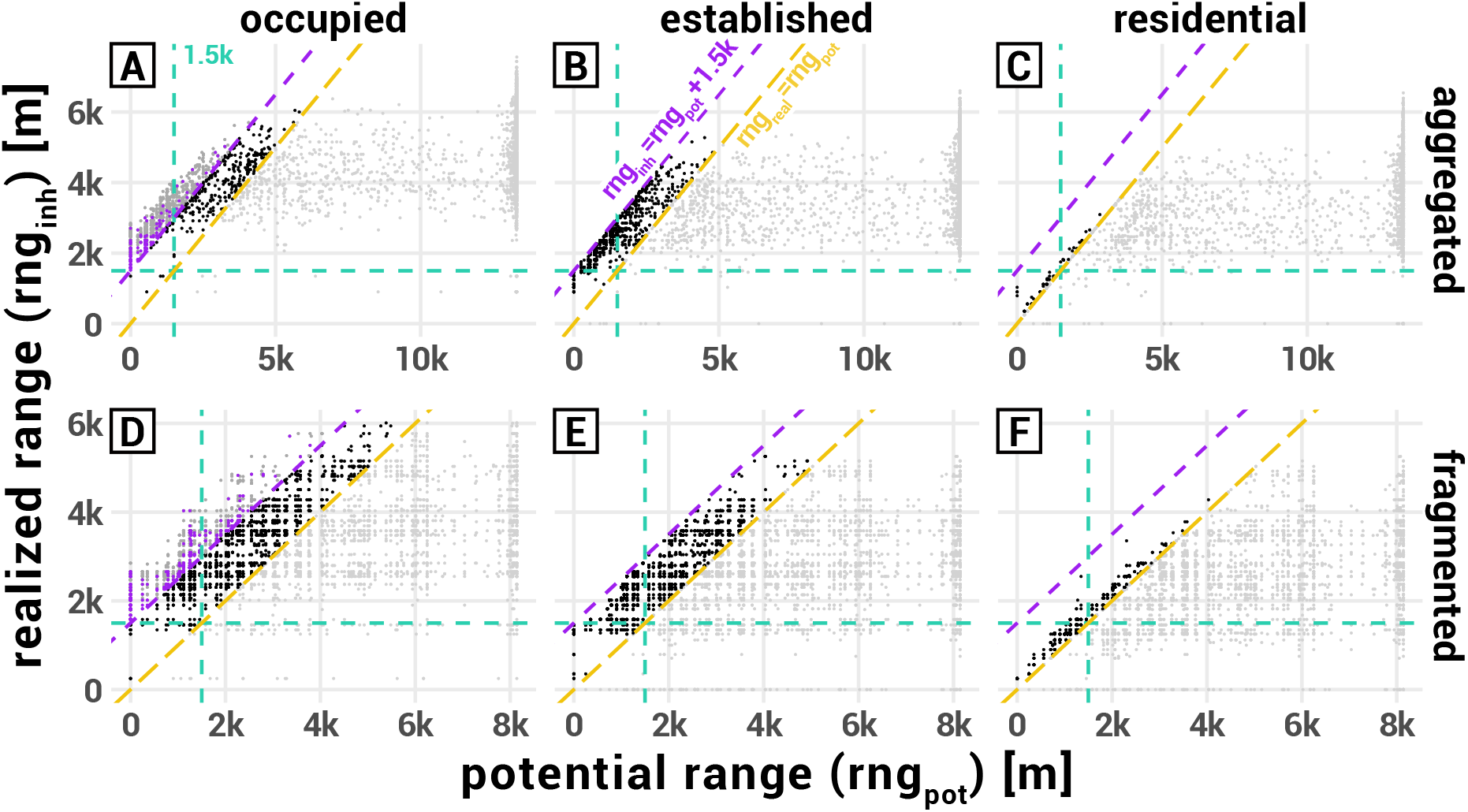
Potential range (*r_pot_*) in meters (x-axis) versus realized range (*r_rel_*) in terms of farthest occupied (A, D), established (B, E) or residential (C, F) habitat during a 60 year simulation run in the aggregated (TOP) or fragmented (BOTTOM) landscape (note different scales in subplots). Each dot represents one replicate run (N=6,000), while the shade of a dot highlights whether *r_real_* remained below (light grey) or within 1,500 m (black) of *r_pot_*, or exceeded it further (dark grey). The diagonal dashed lines mark the respective thresholds (ORANGE: *r_real_* = *r_pot_*, PURPLE: *r_real_* = *r_pot_* + 1,500 *m*). Purple dots additionally mark (occupied) refuges outside the latter threshold. Vertical and horizontal dashed green lines mark the 1,500 m dispersal radius around the habitat of origin. The potential range was only calculated up to respective range under ideal conditions (*p_prot_* = 1.0, RCP8.5), thus the clustering of points at the maximum values of the x-axis.

**Table 2:**
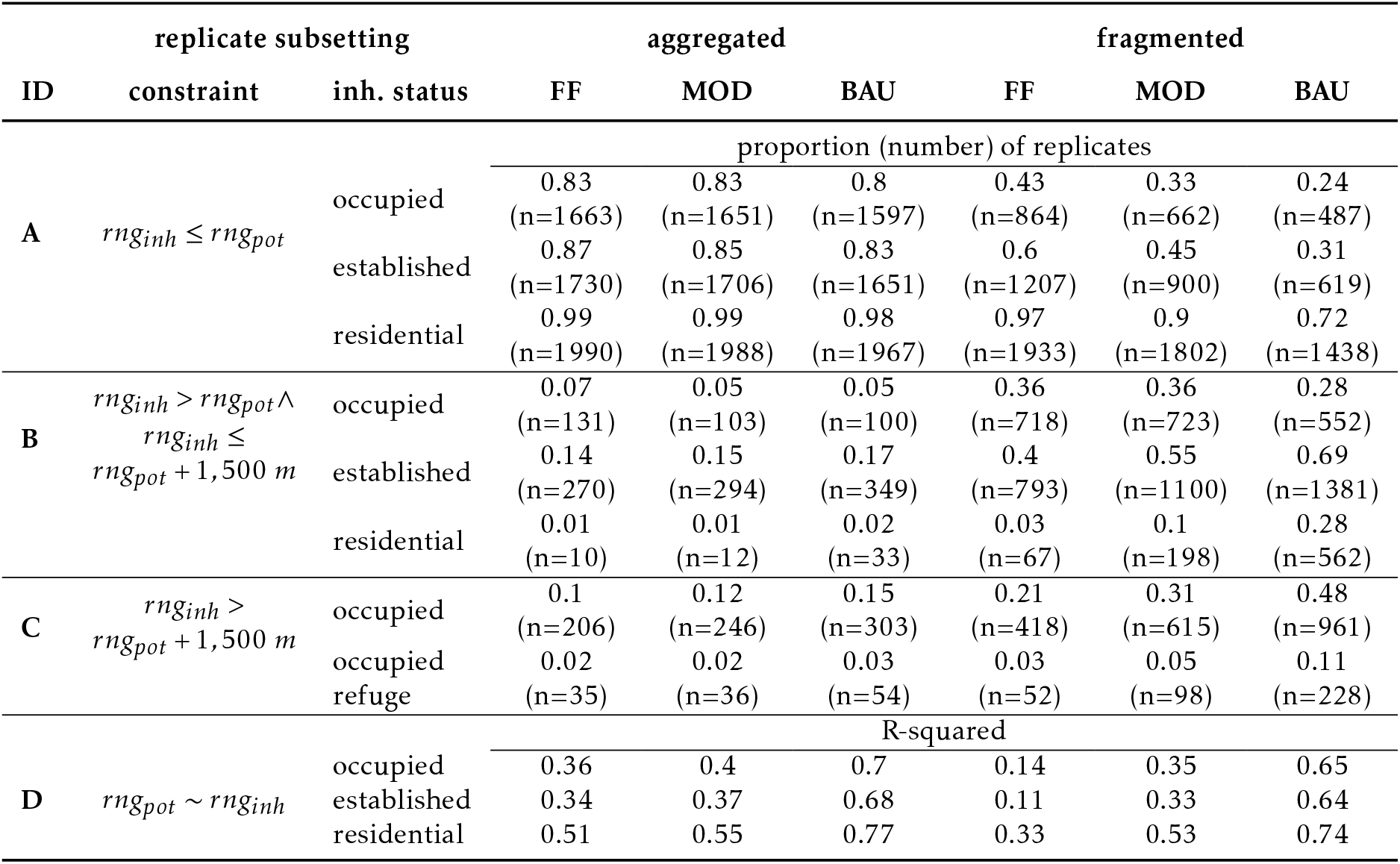
Simulation replicate stats grouped by: (second column) constraint regarding inhabited status dependent range *rng_inh_* compared to potential range *rng_pot_*; (third) inhabited status; and (last six) region and climate change scenario. Upper rows (ID A-C) contain proportion and number of replicates achieving habitats of an inhabited status of certain constraint during simulation run, e.g. with *rng_inh_* remaining within *rng_pot_* (first three rows). Bottom rows (ID D) contain R-squared values for *rng_inh_* dependent on *rng_prot_*

However, the share of protected grasslands influences the pace of the dispersal process within the protected network after a settling phase of five to nine years (Figures 4A-B), depending on region and protected grassland probability. Differences in the residential range depending on the probability of protected habitats slowly become apparent afterwards and grow more significant with advanced simulation time. In the aggregated landscape, it takes on average 29-42 years to find residential habitats outside the dispersal radius of the originating habitat while LMG residents remain within this radius for *p_prot_* ∈ {0.01, 0.02} (fragmented: 39-59 years, *p_prot_* ∈ {0.01,…, 0.06}).

**Figure 4:**
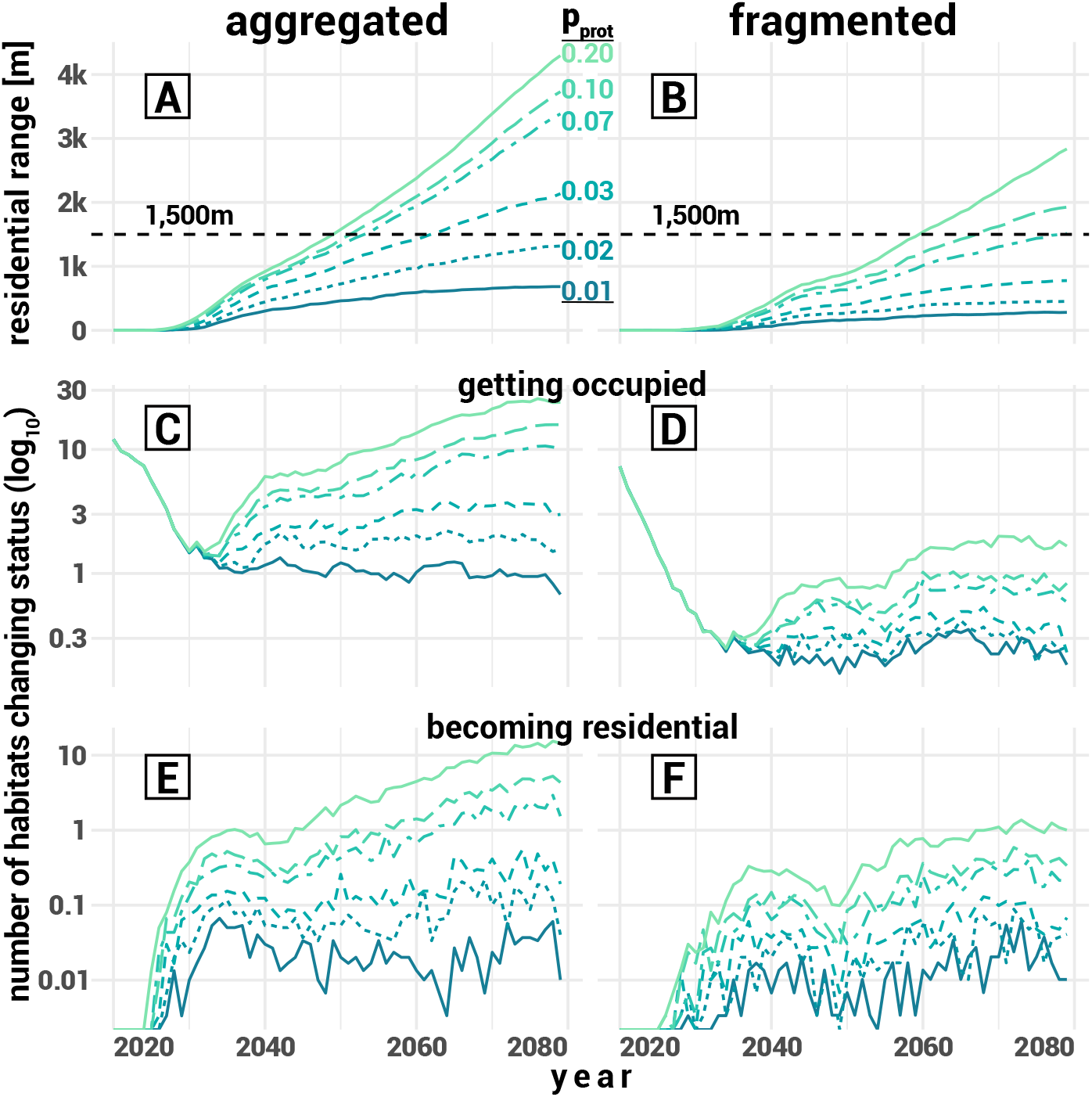
Yearly development (x-axis) of evaluation parameters (y-axis, mean over replicates and climate change scenarios, n=300) by initial region (LEFT: aggregated, RIGHT: fragmented) and (for clarity) selected protected grassland probabilities *p_prot_* (line types and colors). A, B: residential range in meters. C, D: *log*_10_-scaled number of habitats getting occupied for the first time. E, F: *log*_10_-scaled number of habitats becoming residential for the first time. The dashed horizontal line in A and B marks the 1,500 m dispersal radius of the originating location.

The initial delay in achieving residential habitats is explained by the (decreasing number of) newly occupied grasslands during the first years (Figures 4C-D): Singular source of colonization is the population in the originating habitat (thus the decreasing number of new habitats) up to the point where occupied (protected) habitats have grown a large enough population themselves to breed a substantial number of emigrants (Figures 4E-F). Only after that, the maximum residential range is constantly increasing; or diverging for some of the lower *p_prot_* values. The number of grasslands changing their inhabited status enters a fading ‘occupied-residential’ cycle of different extent and slope depending on *p_prot_* and grassland configuration around the habitat of origin (Figures 4C-F). In the fragmented landscape, the rate of yearly occupation-residence remains low for all values of *p_prot_* and never reaches the level of the initial exodus (Figure 4D). This proves to be different in the aggregated landscape, where both rates keep increasing for most of the *p_prot_* values (Figure 4C).

Increasing conservation effort (in terms of *p_prot_*) continuously leads to an extended (residential) range by the end of the simulation in the aggregated landscape (Figure 5, TOP), but the rate of range expansion declines with each additional effort. While for values of *p_prot_* ≤ 0.07 (threshold depending on CCS), every additional percentage point significantly extends the range, above this threshold constantly more effort is required to achieve substantial range expansion. Independent of the CCS, a 3-4 % share of protected grasslands suffices to allow the LMG to become residential in habitats outside the dispersal radius of the originating habitat.

**Figure 5:**
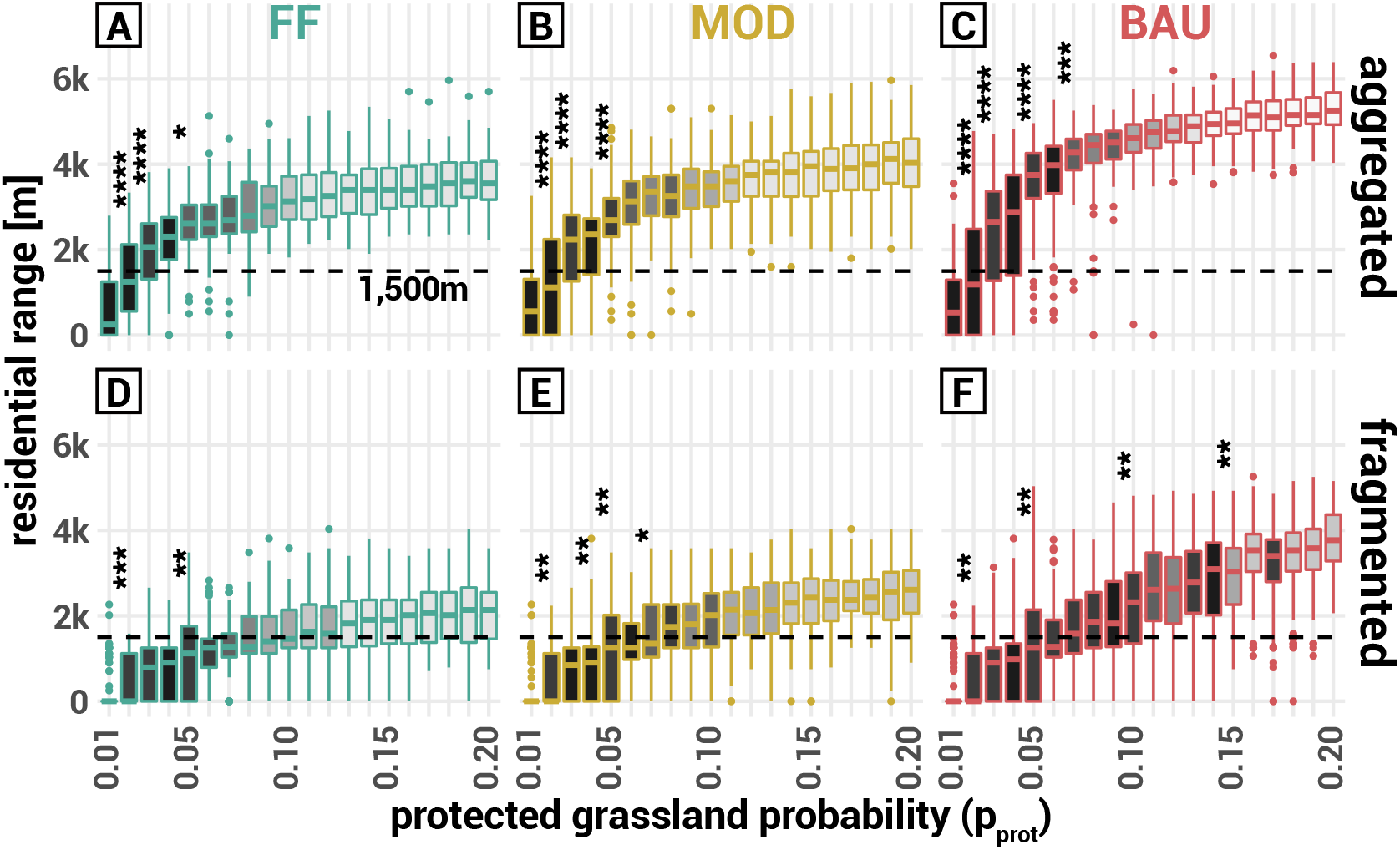
Residential range in meters (y-axis) by the end of the simulation run (2079) in the aggregated (TOP) and fragmented (BOTTOM) landscape depending on protected grassland probability *p_prot_* (x-axis) and CCS (green: FF, RCP2.6; brown: MOD, RCP4.5; pink=BAU, RCP8.5). Grey-scale fill highlights the additional conservation effort (*p_prot_*) required to significantly increase the range, where the darkest grey stands for *p_prot_* + 0.01 and the lightest for *p_prot_* + 0.08 and above. Asterisks mark the P-Value of groups that achieve a significant increase in distance with an additional conservation effort of only 1 % values: **** = *P* ≤ 0.0001, *** = 0.0001 < *P* ≤ 0.001, ** = 0.001 < *P* ≤ 0.01, * = 0.01 < *P* ≤ 0.01

These patterns are different in the fragmented landscape (Figure 5, BOTTOM). Though also here every increase in conservation effort promotes the LMG’s dispersal success to some extent, a share of 7-11 % of protected grasslands (depending on CCS) is required to leave the sphere of the originating habitat after 60 years. Even with higher efforts, the realized residential range remains rather low under mild or moderate climate change. Only under severe climate conditions a high conservation effort allows a substantial dispersal success.

Looking at the population density, there is a highly positive correlation with the value of *p_prot_* (Figure 6). Every additional protected habitat (within potential range of the LMG) aids the species in regionally extending its population. There is only a slightly more significant effect when increasing small *p_prot_* values compared to higher values, but a substantial gain in population density remains when increasing higher values as well.

**Figure 6:**
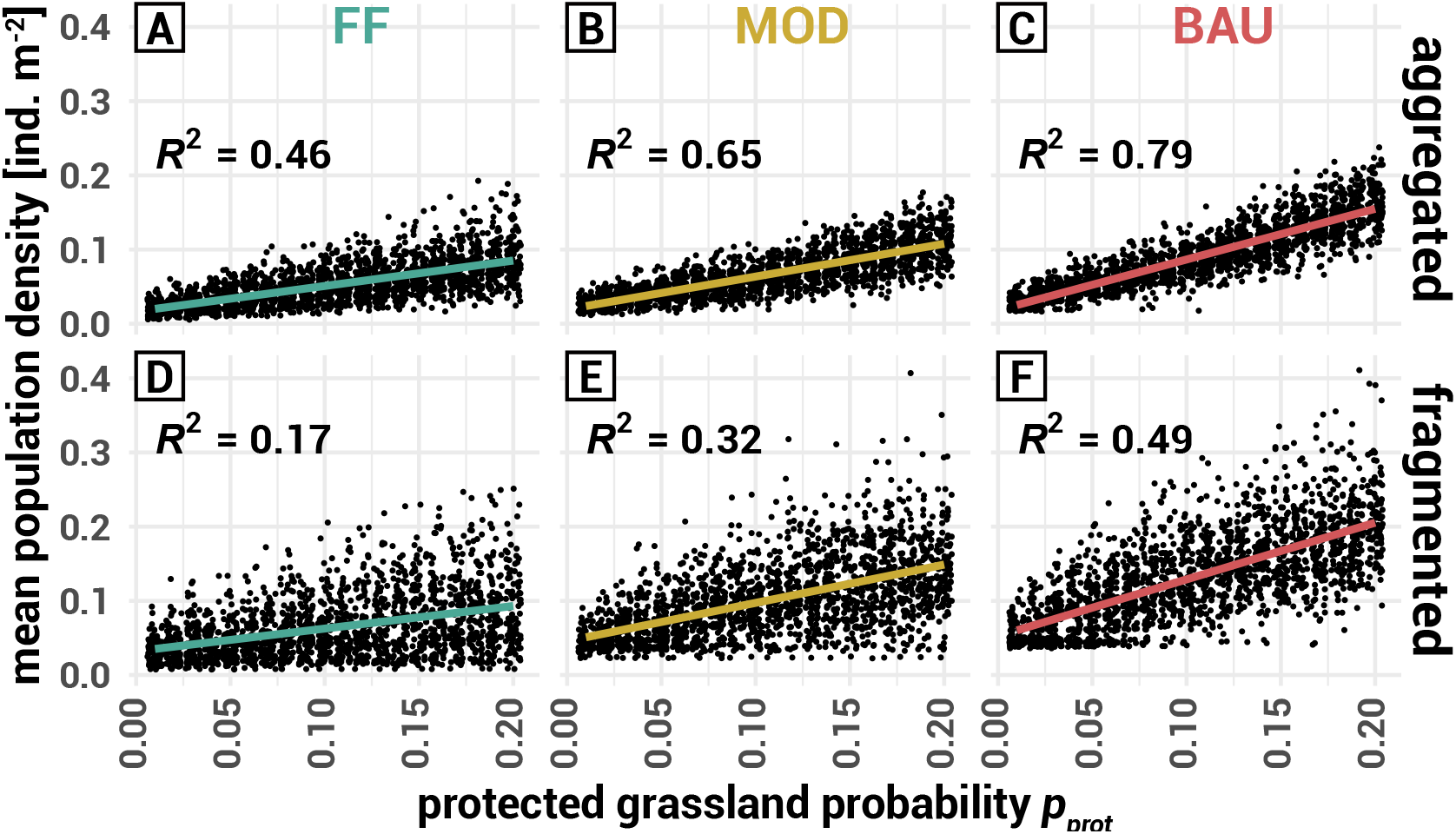
Regional mean population density [*ind. m*^-2^] (y-axis) by *p_prot_* (x-axis) at the end of the simulation run in the aggregated (TOP) and fragmented (BOTTOM) landscape distinguished by climate change scenario (COLUMNS, green: FF, RCP2.6, brown: MOD, RCP4.5, pink: BAU, RCP8.5). Colors highlight the linear trend lines and correlation coefficients.

## 4 Discussion

The following sections discuss the implications for species such as the LMG regarding the effect of poor-quality habitats on weak dispersers (Section 4.1), the relevance of lags in population development for conservation planning (Section 4.2), the increased conservation effort for fragmented landscapes (Section 4.3), and the positive long-term influence limited efforts can have on species development (Section 4.4). Note again the meaning of the inhabited status, as it is important for the interpretation of the results: (1) an *occupied* habitat had an arbitrary small imago density during a year; (2) in an *established* habitat, the overall imago density surpassed a defined threshold but could exclusively consist of immigration; and (3) for determining a residential habitat, immigration is ignored but the density hatched from eggs laid in the preceding year must surpass the same threshold.

### 4.1 Highly Disturbed Habitats Insufficient for Transit of Weak Dispersers

In metapopulation theory (Hanski & Gilpin, 1997), landscapes are usually binary distinguished into terrain suitable and unsuitable for a species. Accounting for a more natural environment, areas that are accessible by a species yet unfavorable for colonization can be considered an additional category of terrain (Wiegand et al., 1999). These so-called poor-quality habitats could facilitate reaching suitable terrain outside the potential dispersal range of a migrating or roving species (Wiegand et al., 2005). At first glance, this facilitation also applies to the results of the present study.

The potential range of the protected network exceeded the dispersal radius of the target species’ originating habitats when protecting only a low percentage of grasslands both in the aggregated and fragmented landscape (Figure 2B-C) Especially in the fragmented landscape, the realized occupied range of the LMG often exceeded the potential range (Table 2C), occasionally with even the closest neighboring refuges outside the species’ dispersal radius (Figure 3, purple dots). The latter clearly indicates that the LMG utilized the whole functional network (of unprotected habitats) during its dispersal process, i.e., passed through poor-quality habitats to reach refuges outside the protected network.

Upon closer evaluation, though, the results indicate that utilizing poor-quality habitats in a highly disturbed environment might not work for species with a short (annual) life cycle and limited dispersal abilities such as the LMG. In contrast to the occasionally clearly exceeding *occupied* range, the realized *established* range always remained within dispersal radius of the closest refuge (Figures 3B and E, black dots) and the *residential* range rarely exceeded the potential range by a few hundred meters (Figures 3C and F, black dots). Especially in the fragmented landscapes, residential populations rather ‘pooled’ on the edge of the potential range (Figure 3F, black dots) suggesting that the environment lacked the means (i.e., habitats suitable enough) for further dispersal.

In most of the cases, the residential range remained well below the potential range during the 60 years (Figure 2C, red marks; Figures 3C and F, light grey dots below orange line; Table 2), although both ranges might eventually align with prolonged simulation time. This alignment is also supported by two factors connected to increasing severity of climate change: (1) more established / residential populations are found outside the potential range (Table 2B), especially in the fragmented landscape; and (2) the correlation between potential range and realized ranges (cf. R-squared values in Table 2D) becomes more pronounced. Thus, an environment better suited for the development of the LMG (i.e., its benefit from climate change, cf. Section 1) promotes realized ranges closer to the potential range.

The observation that grasslands and refuges outside the protected network were occupied but could not sustain an established or residential population indicates that species such as the LMG require a frequent amount of immigration during the initial colonization of a new habitat. Apparently, the connection between refuges via highly disturbed poor-quality habitats of the present simulation setup is insufficient to substantially provide this influx. This observation is in line with the findings of Poniatowski et al. (2018b) that habitat quality is more important for grassland insects (including grasshoppers) than (functional) connectivity between them. Though it is possible that with with increasing number of replicates a remote residential population might occasionally develop, the high number of simulations where the realized residential range remained well below (aggregated) or ‘pooled’ on the edge (fragmented) of the potential range rather suggests that the distance between refuges allowing smooth dispersal lies below the theoretical dispersal radius of the LMG.

This presumed link between refuge distance and smoothness of dispersal is supported by the maximal realized ranges depending on the value of *p_prot_*: In more than 50 % of the cases, relatively low *p_prot_* values build a protected network up to distances unreachable during simulation time (Figure 2C), especially in the aggregated landscape. At the same time, dispersal success further increased (significantly) despite of a relevant increase in potential range (Figures 4, 5, 6). Thus, a higher probability of having (additional) refuges within dispersal radius of other refuges, and therefore reduced distances between them, aids the LMG in occupying more distant refuges within the protected network.

Note that the simulation setup depicts a rather extreme scenario in a landscape of overall highly intensive land use with limited numbers of ideally managed refuges (cf. *p_prot_*). This setup was chosen on purpose to analyze whether it makes sense to implement a (even low-level) diversification of management schedules. As shown in Leins et al. (2021) and Leins et al. (2022), there are management schedules that would support dispersal success and allow reasonable yields at the same time. Applying a heterogeneous (less extreme) setup of management schedules likely could lift the LMG’s restriction on the functional network.

### 4.2 Delay in Establishment Must be Accounted for in the Evaluation of the Conservation Effort

The simulation results show that there is an initial delay in observing established or residential populations aside from the habitat of origin (Figure 4, TOP). With advanced simulation time, observing newly occupied / residential habitats settles into a colonization-residence cycle (Figure 4, MIDDLE / BOTTOM). Hence, the dispersal dynamics of the present model do not follow a continuous diffusion process as postulated in Fick’s laws (Fick, 1855). Foremost the initial lag is a familiar concept in invasion biology (Shigesada & Kawasaki, 1997), and two of its main localized causes, inherent population growth and environmental conditions (Crooks & Soulé, 1999), are also included in the present model. In conservation biology, however, the concept of dispersal lag is, as far as known, not discussed. This is despite the fact that already in classical metapopulation theory (Hanski & Gilpin, 1997), time scales of both local and regional dynamics (e.g. population growth and dispersal processes) are applied and often considered to be interrelated (Drechsler & Wissel, 1997), so that the local dynamics drive the regional dynamics. It has further been reported that in a dynamic landscape, both population growth below a certain threshold and local environmental conditions alter the persistence of a metapopulation compared to the classical theory (Johst et al., 2002). Both the interrelation as well as the altered persistence could possibly result in a delayed dispersal process similar to the present simulation results.

In the dispersal analysis of a range-expanding species, on the other hand, ignoring the effect of local dynamics could result in a qualitative overestimation of dispersal speed, as indicated by the delayed residential range. Therefore, it may be worthwhile for stakeholders in conservation biology to consider the (initial) lag in dispersal during their planning, albeit to enable a species rather than controlling it as intended in invasion biology. Especially for smaller species that are difficult to monitor, such as the LMG, the effect of (initial) lagging phases could be an important factor when assessing newly implemented conservation measures, because their effectiveness might only emerge after a prolonged period of time. While the present study represents a specific case with a singular source of emigration (similar to a species’ reintroduction), the cyclic trend of habitat establishment (Figure 4, BOTTOM) shows that such a delay remains even with advanced simulation time.

The development seen in Figure 4 suggests that newly occupied grasslands require (1) a persistent influx from already established / residential habitats, (2) time to become residential themselves, and (3) a large enough population to be substantial source of emigration. Though in a highly disturbed landscape, as applied in the present study, such a development only works within a network of potentially connected refuges (Figure 4.1), this network does not need to be implemented all at once as indicated by the delayed dispersal process. Depending on the identified species-specific dispersal / establishment rate, conservation planners could initially set up (temporary) refuges within a reasonable radius around known established populations, reevaluate regularly and consecutively add more distant refuges based on the evaluation. Including processes of learning and adapting into conservation planning was suggested before (Grantham et al., 2010) and simulation models such as HiLEG can supply a conceptual basis to support stakeholders in their practical planning.

Achieving the above requirements in a fragmented landscape is much more difficult because of the lower number of available grassland. Even with a high conservation effort there is rarely a single habitat becoming residential per year (Figure 4F), while in an aggregated landscape this is already the case with relatively low effort (Figure 4E).

### 4.3 Required Conservation Effort Significantly Higher for Fragmented Landscapes and Minor Climate Change

In general, the simulation results indicate that, for a species benefiting from climate change such as the LMG, required conservation effort would be the lowest under severe climate conditions and in an aggregated landscape (Figures 4, 5, 6). While the positive response of the LMG regarding climate change was already shown in previous studies (Leins et al., 2021; Poniatowski et al., 2018a; Trautner & Hermann, 2008), it is reasonable to focus conservation planning on conditions more challenging for the species. First, because the United Nations remains committed to achieving the 1.5 °C goal (Paris Agreement, 2016), i.e., a less beneficial scenario for the LMG. Second, because the effect of measures can only be observed after some years (cf. Chapter 4.2), focusing a broader conservation effort on short-term gain could be convenient (Wilson et al., 2006).

However, it is questionable, whether it makes sense to expend any effort on protecting grasslands in a fragmented landscape. Allowing the LMG to achieve self-sustaining populations in distances rather close to the origin might already take decades and require a high number of protected grasslands (Figures 4B), especially under less severe climate conditions (Figures 5D-E).

Even when considering the trend that every additionally protected grassland helps increasing the population density (Figure 6), it does not necessarily apply to the less severe climate conditions in the fragmented landscape. The correlation between the protected grassland probability *p_prot_* and population density is weak under minor climate conditions (Figure 6D, *R*^2^ = 0.17) and low under medium conditions (Figure 6E, *R*^2^ = 0.32), so it is not guarantied that implementing additional protected grasslands would have a substantial effect.

In scenarios of the aggregated landscape, the effect of conservation effort on population development is higher. Increasing the probability of protected grassland quickly reflects in success of occupying, establishing or residing in more distant habitats (Figures 4A, 5A-C) and correlates better with population density (Figures 6A-C), especially in the more severe CCS. Focusing conservation planning on LMG populations present in aggregated landscapes might thus prove more sustainable and effective.

### 4.4 Slightly Increasing Low Conservation Effort Can Have Positive Long-term Effect

In cultivated grasslands, it might prove difficult to provide incentives for farmers or other stakeholders in order to implement suitable measures. A recent review by Nguyen et al. (2022) highlights, for instance, that despite the general recognition to preferably apply conservation measures at a landscape-scale, there are few real-world examples of such implementations.

However, as discussed before, every additional effort in protecting grasslands could support the population development of the LMG, especially in the aggregated landscape (Figures 5, 6, TOP). Here, the simulation results show that protecting a low percentage of 1 – 3 % of grasslands already has a notable effect on the dispersal success or rather allows self-sustaining populations aside from the habitat of origin (Figures 5A-C). More importantly, each additional percentage point in the low single digits allows a significant expansion of the realized residential range by the end of the simulation run (cf. black boxes marked with asterisks in Figure 5).

While increasing the conservation effort further still has an effect on both realized ranges and - as discussed before - the population density (Figure 6), it requires a constantly larger effort to achieve a significant response in range (cf. grey scale of boxes in Figure 5). Therefore, it can be sufficient to achieve a small number (1 – 3 %) of protected grasslands (in vicinity of an already protected population) and smartly increasing the number over time (cf. 4.2) up to a reasonable threshold (5 – 8 %). Above this threshold, additional effort would not be in vain, but no longer have a strong effect.

Regarding the review mentioned above, the present simulation results offer the prospect that for species such as the LMG, even localized measures could have a positive effect, although coordinated conservation effort at the landscape-scale would be more beneficial on the long run.

## 5 Conclusion

In a highly disturbed or intensively managed environment, protected grasslands might only have a limited positive effect on species with low dispersal ability and a rather short life cycle such as the large marsh grasshopper (LMG). On a regional scale, a population remains restricted to refuges within its (in)direct dispersal range and cannot make use of intensively managed grassland as transit habitats to sustainably establish in refuges outside a network of protected habitats that are (in)directly connected by the species’ maximum dispersal radius. Within a reasonable period of time, populations might in fact often only establish in distances well below the potential range of this protected network. Placing refuges closer together as required by a species’ dispersal radius can notably aid it in dispersing farther and establish a robust core population. This positive effect is particularly evident when, in terms of dispersal radius, larger distances between refuges are reduced and becomes less striking when refuges were already closer together. Therefore, it can be more beneficial to create some (additional) neighboring refuges in an area with none or few protected grasslands than to do so in an area with already nearby refuges.

When implementing such refuges, it can be of importance to consider the potential delay in local population development of species with similar traits as the LMG. It can take several years to have a visible effect of the conservation effort. First, it may require some time for a species to find a suitable habitat, and second, the species needs to develop a population size large enough to measure during a survey. Such a delay on the spatial edge of a range-expanding dispersal process can lead to a cyclic colonization behavior that can easily be overlooked. Stakeholders should keep the delay in mind when assessing the effectiveness of their conservation measures for species difficult to monitor.

Focusing the conservation efforts on an aggregated landscape is much more promising than on a fragmented environment. The probability of achieving a protected habitat in range of another is higher within an agglomeration of grasslands. Furthermore, despite the fact that the intensively managed grasslands do not aid in dispersing to refuges outside the protected network, they could still function as temporary habitats and thus contribute to the overall population development. This is rarely the case in a fragmented environment.

In general, the results show that even in a rather extreme setup of a intensively managed environment with the occasional refuge, species of limited dispersal ability could establish to some extent and range. Implementing a more heterogeneous setup of land use management (i.e., schedules with different levels of negative impact on a target species’ development) should allow a species to expand even outside the range of locations considered protected habitats. Previous studies showed that such intermediate schedules exist. HiLEG can be used to explore such a heterogeneous setup to identify management schedules that promote a target (grassland) species in thriving regionally.

## Supporting information

ODD model description

## Footnotes

1 GitLab repository of HiLEG release version v1.5: https://git.ufz.de/leins/hileg/-/tree/v1.5

2 Output data (spatially aggregated region): https://www.ufz.de/record/dmp/archive/12742/en/

3 Output data (spatially fragmented region): https://www.ufz.de/record/dmp/archive/12741/en/

4 Evaluation data: https://www.ufz.de/record/dmp/archive/12743/en/

