## Supplementary material for "Finding the right balance of conservation effort in cultivated grasslands: A modelling study on protecting dispersers in a climatically changing and anthropogenically disturbed environment": ODD model description

### Supplement S1: ODD model description of the *Hi*gh-resolution *L*arge *E*nvironmental *G*radient simulation model

The model description below follows the ODD (Overview, Design concepts, Details) protocol (Grimm et al., 2006; Grimm et al., 2020). It includes the original model description (M1 and black text) used to study the effects of land use and climate change on spatially stationary populations of the large marsh grasshopper in Northwest Germany (Leins et al., 2021); it describes extensions made to the original model (M2 and green text) to explore the additional effects on the species when adding dispersal in an environment of higher spatial resolution (Leins et al., 2022); and it incorporates the description of changes made (M3 and pink text) to study the implications of varying conservation effort for the present work.

#### 1 Purpose and Patterns

M1: The *PURPOSE* of the model HiLEG (*Hi*gh-resolution *L*arge *E*nvironmental *G*radient) is to answer the following questions regarding populations of the large marsh grasshopper (LMG) Stethophyma grossum (Linné 1758) in Northwest Germany: (1) How do population density and viability shift regionally, given different climate change scenarios? (2) Which mowing schedule has the least negative impact on the overall population density and viability in the study region? (3) Does the mowing impact severity depend on the spatial location (with its specific climate)?

The empirical patterns used to ensure that the model is realistic enough for its purpose are observed features of the life cycle and their sensitivity to environmental conditions, which were taken from literature. These patterns were used for the model's design. Model output in terms of population structure, densities and persistence were not compared to data, as such data are sparse. Therefore, all model predictions are relative, not absolute. The model was implemented in C++. The source code of the model implementation and the input files used for the simulations runs are available via a GitLab repository[1].

M2: The *PURPOSE* of the model extension is to study the additional effect of dispersal on the LMG in a North German environment of realistic grassland distribution with higher spatial resolution to answer the following questions: (1) Are there (regional) differences in dispersal success depending on climate change scenario? (2) Is the success of dispersal additionally

[1] HiLEG GitLab repository: `git.ufz.de/leins/hileg`

affected by spatial patterns such as grassland cover? (3) Can dispersal compensate for otherwise detrimental grassland mowing?

The species' dispersal metrics were taken from literature and known LMG habitats (Leins et al. (2022), Figure 2B, orange circles) adapted from survey data[2] gathered in the years 2000 to 2016, which were used to analyze some implications of regional effects. Other measures of dispersal success are relative, not absolute, due to a lack of relevant data.

M3: *PURPOSE* of the second extension of the HiLEG model is to study which are the implications of restricted, heterogeneous conservation effort targeting the LMG addressed by these research questions: (1) How does the relative effort in conservation-oriented grassland management affect the population development of a species with limited dispersal ability? (2) Are there time-critical factors that are worth considering for conservation planning in a climatically changing environment? (3) Does the conservation effort required to meet a conservation target differ depending on the spatial landscape structure?

Two known LMG populations on the edge of unpopulated grasslands (taken from survey data[2]) were contemplated to examine the effect of spatial composition (aggregated, fragmented) on the dispersal success of the range-expanding species in a disturbed and changing environment. Timing of land use (mowing) was coupled to the start of the vegetation period as suggest by Gerling et al. (2020).

#### 2 Entities, State Variables and Scales

M1: The model has the following entities: *Grid Cells* (defining environmental conditions) and *Population* per *Grid Cell* comprised of *Life Stages* which are comprised of age-distinguished *Cohorts*. *Flows* are auxiliary entities that manage the *density transfer* between *Life Stages* or their *loss* through mortality.

M2: Instead of *Grid Cells* the model extension has two separate entities *Climate Cells* (defining large scale climate conditions in a $12 \times 12\ km^2$ region) and *Grassland Cells* (defining environmental conditions, e.g. interpolated climate values, on a scale of $250 \times 250\ m^2$). A *Grassland Cell* contains an initially empty *Population* entity and is considered *inhabited* if the *Population* has a non-zero *density*. Otherwise it is considered *uninhabited*. The *Flow* entities additionally connect *Grassland Cells* and handle the *density transfer* between their *Populations* during dispersal.

M1: The LMG develops through three main *Life Stages* during a year (cf. Leins et al. (2021), Section 2.2). Following Ingrisch (1983) and Wingerden et al. (1991), we divided the egg / embryo stage into pre-diapause, diapause and post-diapause development (called embryo hereafter) to account for the clutch's different susceptibility to climate conditions in autumn, winter and spring. This subdivision yields five *Life Stages*: (1) pre-diapause, (2) diapause, (3) embryo, (4) larva, (5) imago. Stages (1) to (3) occur below ground, stages (4) and (5) above ground. Furthermore, stages (2) and (4) can have multiple *Cohorts*, to allow survival over several years in case of conditions during winter that are unsuitable for development, and to account for different temperature-driven development speed depending on hatching date.

M2: Dispersal only occurs between the *Populations'* imago *Life Stages* of *Grassland Cells* within a defined neighborhood.

[2]Provided by Landesamt für Landwirtschaft, Umwelt und ländliche Räume via our project partner Stiftung Naturschutz Schleswig-Holstein

M1: Tables S1.1 and S1.2 provide an overview of the model's life cycle and spatial entities as well as their state variables. The *Population* consists of several *Life Stages* and is characterized by the *coordinates* of its *Grid Cell* and a *minimum density* [individuals $m^{-2}$]. The *Population*'s *density* and *aboveground density* [both in individuals $m^{-2}$] are calculated from its *Life Stages' densities*. A *Life Stage* has a *name*, consists of one or more *Cohorts* and supplies a *maximum age* [days] for each of those *Cohorts*. *Cohorts* exceeding the *maximum age* or falling below the *minimum density* are considered extinct. Furthermore, a *Life Stage* has a *flag* informing whether it occurs below or above ground. The *Life Stages' density* [individuals $m^{-2}$] is calculated from its *Cohorts' densities* and the amount of *gain* [individuals $m^{-2}$] from the incoming *transfer Flow* of its preceding *Life Stage*.

*Cohorts* are distinguished by an *ID* and have an *age* [days] and a *density* [individuals $m^{-2}$]. They have a *development progress* given as a ratio $\in [0,1]$ that defines if and how *density* is transferred to subsequent *Life stages*. The *development progress* is used in two different ways depending on the external *Influence* (Section 7.1) associated with the *transfer Flow*: (1) it is directly affected by external *Influences* and contributes to a *Flow*'s *density transfer* to the subsequent *Life Stage* if it reaches a value of 1 (Section 7.4); or (2) it is used within stochastic *Influences* to determine the probability and extent of contribution to the *flow rate* (Section 7.1, Binominal Climate, M2: renamed from Factor; Algorithm S1.2). Without external *Influences*, the *development progress* defaults to a value of 1.

M1: The auxiliary entity *Flow* was introduced to ease the implementation of the model. Each *Flow* is characterized by the *life stage of origin* and *life stage of destination* (empty for *mortality Flow*), a static per capita *base flow rate* [$day^{-1}$] and a *total flow amount* [individuals $m^{-2}$]. For each *Cohort* in its *life stage of origin* it has a per capita *dynamic flow rate* [$day^{-1}$] and a *current flow amount* [individuals $m^{-2}$]. These *flow rates* are calculated using external *Influences* (Section 7.2). The different *types* of *Flows* define the different flow processes: *trans* (transferring *density* to a subsequent stage: $1 \rightarrow 2$, $2 \rightarrow 3$, $3 \rightarrow 4$, $4 \rightarrow 5$); *repr* (reproduction from imago (no loss) to pre-diapause stage; $5 \rightarrow 1$). Additionally, all five *Life Stages* and their *Cohorts* lose *density* through *mortality* (*Flow type mort*).

M2: A fourth *Flow type disp* (dispersal) is introduced defining the *density transfer* from a *Life Stage* to the same *Life Stage* (here, imago only) of a neighboring *Population*. Potential *density loss* during dispersal of the imago *Life Stage* is handled by an additional *mortality Flow*.

M1: The *Grid Cells* comprising the environment are characterized by their *coordinate* (cell indexes), *carrying capacity* (maximum number of aboveground population that can be sustained), daily climate conditions (*temperature*, *humidity*, *contact water*) and land use schedule (*mowing day*).

M2: *Grid Cells* are replaced by the two entities *Climate Cell* and *Grassland Cell* with the indexes of the former and the geometric center of the latter belonging to the same Cartesian coordinate systems. In this coordinate system, *x-coordinates* increase from West to East and *y-coordinates* from North to South. *Climate Cells* have a unique *ID* and contain the daily climate conditions in the original spatial resolution. *Grassland Cells* take the *carrying capacity* and a *mowing schedule* while adapting the climate conditions of up to four adjacent *Climate Cells* to calculate the local climate conditions using bilinear interpolation. Both the *mowing schedules* and the bilinear interpolation will be described in more detail below.

M1: The model uses daily time steps for updating the model's states and process. This time scale also reflects the sampling of the climate data. However, the single possible mowing event per year is considered on a weekly basis. To account for this weekly frequency, a year has 364 days by definition, resulting in exactly 52 full calendar weeks $year^{-1}$. Input data (i.e.,

**Table S1.1:** Overview of the model's life cycle entities (first column) and their state variables (second column). The text in parentheses of columns one and two represents the entity's or state variable's symbol when used, e.g., in equations. The third column gives the (initial) value(s) of the state variables, the fourth column gives their units (if any). In the fifth column, a brief description of the state variable is provided. The indexes for location and time step that distinguish entities and their dynamically changing states are implied and not explicitly specified in the identifier. The parameter $A_{hab} = 62,500\ m^2$ is the habitat size modeled for a Population.

| Entity (symbol) | State Variable (symbol) | Value(s) | Unit | Description |
|---|---|---|---|---|
| Population ($P$) | coordinate ($coord_{x,y}$) | $x, y \in [1, 36]$ | | Index of rotated pole grid coordinates (see Leins et al., 2021) |
| | set of Life Stages ($P^{stages}$) | $\{S^{pre}, S^{dia}, S^{emb}, S^{lar}, S^{ima}\}$ | | Distinguished Life Stages of the target species |
| | set of Flows ($P^{flows}_{type}$) | | | A set of all Flows associated with this Population distinguished by their flow type (see below) |
| | density ($dens^P$) | $\sum\{dens^{pre}, dens^{dia}, dens^{emb}, dens^{lar}, dens^{ima}\}$ | $ind.\ m^{-2}$ | Summed Life Stage densities |
| | aboveground density ($dens^P_{above}$) | $\sum\{dens^{lar}, dens^{ima}\}$ | $ind.\ m^{-2}$ | Summed density of the aboveground Life Stages |
| | minimum density ($dens_{min}$) | $\frac{1}{A_{hab}}$ | $ind.\ m^{-2}$ | Minimum density of one individual per habit |
| Life Stage ($S^{name}$) | name | $\in \{pre, dia, emb, lar, ima\}$ | | Name of the distinguished Life Stages |
| | set of Cohorts ($S^{name}_{cohorts}$) | $\{C^m, \ldots, C^n\}, \{m, n\} \in \mathbb{N}$ | | Distinguished Cohorts associated with the Life Stage |
| | density ($dens^{name}$) | $\sum\{dens^m, \ldots, dens^n\}, \{m, n\} \in \mathbb{N}$ | $ind.\ m^{-2}$ | Summed densities of the associated Cohorts |
| | gain ($gain^{name}$) | | $ind.\ m^{-2}$ | Summed total flow amount of the incoming Flows (Sections 7.3, 7.4) |
| | aboveground flag ($above^{name}$) | $\in \{TRUE, FALSE\}$ | | Boolean flag defining whether the stage occurs above ground |
| | maximum age ($dens^{name}_{max}$) | $\in \{210, 1700, 120, 90, 120\}$ | days | Maximum age of the associated Cohorts |
| Cohort ($C^{name}$) | ID | $\in \mathbb{N}$ | | Unique Cohort identifier |
| | density ($dens^{ID}$) | | $ind.\ m^{-2}$ | The Cohort's individual density |
| | age ($age^{ID}$) | | days | Age in days since Cohort creation |
| | development progress ($prog^{name}_{ID}$) | $\in [0, 1]$ | | The ratio of development compared to full development |
| Flow ($F^{name}_{type}$) | type | $\in \{trans, repr, mort, disp\}$ | | Defines how the Flow is processed |
| | life stage of origin ($orig$) | $\in \{S^{pre}, S^{dia}, S^{emb}, S^{lar}, S^{ima}\}$ | | Life Stage used to determine the amount of flow |
| | life stage of destination ($dest$) | $\begin{cases} \text{mort}, & \text{if type=mort} \\ \in \{S^{pre}, S^{dia}, S^{emb}, S^{lar}, S^{ima}\}, & \text{otherwise} \end{cases}$ | | Life Stage receiving the amount of flow, or the amount of mortality |
| | set of influences ($I^{name}_{type}$) | | | Environmental drivers associated with this Flow |
| | base flow rate ($rate^{name}_{type}$) | | $day^{-1}$ | Daily per capita base flow rate |
| | dynamic flow rate ($dyn^{ID}_{type}$) | | $day^{-1}$ | Daily per capita flow rate per Cohort in the life stage of origin |
| | current flow amount ($amount^{ID}_{type}$) | $dens^{ID} \times dyn^{ID}_{type} \times day$ | $ind.\ m^{-2}$ | Amount of density flow over one day |
| | total flow amount ($amount^{name}_{type}$) | $\sum\{amount^m_{type}, \ldots, amount^n_{type}\}, \{m, n\} \in \mathbb{N}$ | $ind.\ m^{-2}$ | Summed amount (per Cohort) of density flow over one day |

**Abbreviations: above=aboveground, C=Cohort, coord=coordinate, dens=density, dest=destination, dev=development, dia=diapause, disp=dispersal, dyn=dynamic, emb=embryo, F=Flow, hab=habitat, ID=Cohort identifier, ima=imago, ind=individuals, lar=larva, m=meter, max=maximum, min=minimum, mort=mortality, orig=origin, P=Population, pre=prediapause, prog=progress, repr=reproduction, S=Life Stage, t=time step, trans=transfer**

**Table S1.2:** Overview of the model's spatial entities (first column) and their state variables (second column). The text in parentheses of columns one and two represents the entity's or state variable's symbol when used, e.g., in equations. The third column gives the (initial) value(s) of the state variables, the fourth column gives their units (if any). In the fifth column, a brief description of the state variable is provided. The indexes for location and time step that distinguish entities and their dynamically changing states are implied and not explicitly specified in the identifier.

| **Entity (symbol)** | **State Variable (symbol)** | **Value(s)** | **Unit** | **Description** |
|---|---|---|---|---|
| Climate Cell ($\Omega$) | ID | $\in [1, 107] \subset \mathbb{N}$ | | Unique identifier of Climate Cells in Schleswig-Holstein |
| | $center_{x,y}$ | $\{x, y\} \in \mathbb{N}$ | | Geometric center in coordinate system of Grassland Cells |
| | temperature ($\omega_{ts}$) | | °C | Local surface temperature |
| | humidity ($\omega_{rhug}$) | | % | Local relative humidity in the upper 2 cm of the ground |
| | contact water ($\omega_{cw}$) | | $kg\ m^{-2}$ | Amount of water in the upper 2 cm of the ground |
| Grassland Cell ($G$) | coordinate ($coord_{x,y}$) | $\{x, y\} \in \mathbb{N}$ | | Index in Cartesian coordinate system |
| | carrying capacity ($cap_{above}$) | 25 | $ind.\ m^{-2}$ | Maximum aboveground density |
| | climate value(s) ($\omega_{clim}$) | | | Calculated by weighing resp. values of adjacent Climate Cells |
| | mowing schedule ($T_{mow}$) | See Table S1.4 | days | Occurrence days of mowing events |
| | start of vegetation period ($t_{veg}$) | | day | Day on which the yearly sum of surface temperature $\omega_{ts}$ reaches 200.0 °C (see Eqn. S1.28) |

**Abbreviations: above=aboveground, cap=capacity, clim=climate, coord=coordinate, cw=contact water, G=Grassland Cell, ID=Climate Cell identifier, ind=individuals, kg=kilogram, m=meter, mow=mowing, rhug=relative humidity upper ground, t=time step, temp=temperature, ts=surface temperature, veg=vegetation**

climate time series) is either cropped or expanded accordingly. Simulations were run for 20 years (7280 time steps) or stopped earlier in case all *Cohorts* of the *Population* became extinct.

M2: For each year of the climate data, February 29$^{th}$ (if exists) and December 31$^{st}$ are omitted to achieve 364 days. A simulation run takes 21,840 time steps (60 years) starting on January 1$^{st}$ 2020 and ending on December 30$^{th}$ 2079. In the case of premature extinction of all *Populations*, simulations stop earlier.

M1: The environment, or study region, comprises 1296 ($36 \times 36$) cells, each having an area of 144 $km^2$ ($12 \times 12$ $km^2$), which corresponds to the resolution of the climate input data (Section 6). The integer *Grid Cell* indexes (1-36) are mapped to rotated pole grid coordinates (Leins et al. (2021), Figure 1). 968 *Grid Cells* are terrestrial and therefore belong to the model domain. Within a *Grid Cell* a single habitat is simulated and represents a squared virtual grassland plot with the size of 6.25 ha ($250 \times 250$ $m^2$). *Grid Cells* and hence habitats are not connected, i.e., there is no exchange of individuals. If populations become extinct, there is no recolonization.

M2: The study region is the German federal state of Schleswig-Holstein (SH) consisting of a 107 terrestrial *Climate Cell* subset of the original data and 72,969 *Grassland Cells* representing the state's actual grassland area that were retrieved using the software DSS-Ecopay (Mewes et al., 2012; Sturm et al., 2018). Each *Grassland Cell* by definition has a size of 6.25 ha ($250 \times 250$ $m^2$) that reflects the spatial resolution of the data quite well. *Grassland Cells* within a radius of 1,500 m are connected by the dispersal process of the imago *Life Stage*. In some cases, there are additional connections outside this radius representing long distance dispersal (LDD). The connections and dispersal processes will be described below.

M3: Two sub regions of the grasslands described above function as point of origin for the simulation of a range-expanding species, both containing a known LMG population on the edge of unpopulated territory. The regions are of different spatial composition (cf. Leins (2022), Figure 2B), where the northern one is spatially fragmented in terms of grassland cover around the known population and the southern one has rather aggregated grasslands. Again, *Grassland Cells* are connected by the dispersal process in a radius of $1,500\ m$, but LDD is ignored to focus the analysis on the actual grassland composition.

#### 3 Process Overview and Scheduling

M1: In each time step and for every *Grid Cell* (M2: *inhabited Grassland Cell*), four main blocks of *PROCESSES* are executed: 'Update environmental drivers', 'Flow update', 'Life Stage update', and 'Cohort update'. The first three blocks are SCHEDULED one after the other, while 'Cohort update' is executed as a submodel of 'Life Stage update'. Algorithm S1.1 gives an overview of this top level scheduling. During 'Flow update' the *flow rates* and *amounts* are calculated using the *Flow*'s *life stage of origin* while depending on its *type* and specified external *Influences* (submodel 'Update environmental drivers').

M2: The *PROCESS* 'Bilinear climate interpolation' is executed each time step as a submodel of 'Update environmental drivers'. If the *life stage of destination* of a non-zero (in terms of *flow amount*) dispersal *Flow* belongs to the *Population* of an *uninhabited Grassland Cell*, the *Population* is not empty anymore and the cell thus rendered *inhabited*. In this case, the submodel 'Dispersal setup' is executed to find all *Grassland Cells* considered neighbors of the *Population* and establish a dispersal connection to each of them by creating a respective *Flow* of *type disp*.

M3: The *PROCESS* 'Start of vegetation period' is executed executed each time step as a submodel of 'Update environmental drivers' right after submodel 'Bilinear climate interpolation'.

M1: The 'Life Stage update' first handles creation (input from *Flows*) and lastly deletion (*density* falling below *minimum density*) of its *Cohorts*. In between *Cohort* creation and deletion, the submodel 'Cohort update' is executed.

**Algorithm S1.1** Main process overview. Processes executed at each inhabited Grassland Cell and for each time step. The processes 'Bilinear climate interpolation' and 'Cohort Update' are executed as submodel of 'Update environmental drivers' and 'Life Stage update', respectively.

```
for all Inhabited Grassland Cells / Populations do
    RUN SUBMODEL('Update environmental drivers')
    for all Originating Flows do
        RUN SUBMODEL('Flow update')
        if [Flow_type = 'dispersal' & Flow_amount > 0 & Population^target_density = 0] then
            Grassland Cell^target_state ← 'inhabited'
        end if
    end for
end for
for all Inhabited Grassland Cells / Populations do
    Population_density ← 0
    for all Life Stages do
        RUN SUBMODEL('Life Stage update')
        Population_density ← Population_density + Life Stage_density
    end for
    if Population_density < min_density then
        Grassland Cell^target_state ← 'uninhabited'
    end if
end for
```

### 4 Design Concepts

*Basic Principles:*

M1: The model uses viability, i.e., the ability of small populations to persist, as a comparative metric to identify grassland *mowing schedules* that are compatible with the projected climate conditions in a certain region. Development and mortality of the different *Life Stages* are driven by environmental factors. For this, relationships were imposed that reflect the available empirical knowledge and data.

M2: The extension uses the same comparative metrics and environmental drivers as the original study. Additionally, it applies density-independent dispersal using a fat-tailed dispersal kernel that changes depending on regional grassland cover. Relevant dispersal parameters were adapted from empirical studies of the target species.

*Emergence:*

M1: The relationships describing the life cycle are imposed and not emergent from first principles, such as energy budgets or adaptive decision making.

M2: The dispersal metrics are imposed by the parameter definitions such as dispersal radius.

*Interaction:*

M1: The model does not include direct interaction within or among different *Life Stages*. Indirect interactions are included by assuming density dependence of the mortality of larvae and imagines.

M2: Neighboring *Populations* of *inhabited Grassland Cells* indirectly interact through a dispersal process that depends on the grassland cover surrounding the originating cell.

*Stochasticity:*

M1: Transfer from embryo to larval ($F^{emb}_{trans}$, Table S1.5) and larval to imago ($F^{lar}_{trans}$) *Life Stage* is stochastically drawn from a binominal distribution (Section 7.1). The probability is influenced by temperature, *Cohort density* and *development progress*.

M2: Dispersal mortality as well as the actual dispersal from a *Population*'s imago *Life Stage* to the imago *Life Stages* of its connected *Grassland Cells* is determined stochastically using a density dependent binomial distribution. The *dispersal probability* for each connection is calculated using a predefined *base dispersal rate*, a distance dependent *dispersal preference*, a *probability* of finding the connected neighbor and a *survival probability*, where the latter two depend on grassland cover and distance between both *Grassland Cells*. The probability of dispersal mortality is derived from the summed dispersal probabilities.

### 5 Initialization

M1: The model is initialized with a *starting date*, *duration*, *mowing day* and *climate change scenario* (CCS). Additionally, each *Population* receives an *initial density* per *Life Stage*. The *carrying capacity* [individuals $m^2$] for the aboveground population is assumed to be the same for all *Grid Cells* to reduce the number of confounding factors in the model analysis. As the climate data only offers a single projection per RCP scenario (cf. Leins et al. (2021), Section 2.3) and location, we resampled the time series per replicate run (*seed*): for a simulation period of 20 years, we used a time frame of 30 years (simulation period ±5 years) and reordered the years by sampling with replacement. This resampling is feasible since there are no significant trends in the relevant climate parameters within such a period (K. Keuler, pers. comm.). Simulation period 2060-79 has just 26 years, because climate data was only available up to the year 2080. A list of the sampled years per *seed* and simulation period is supplied in Leins et al. (2021), Supplement S2. The initial settings of the model are listed in Table S1.3.

M2: Every simulation starts on 01 January 2020 and runs for 60 years (21,840 time steps). A run is initialized with one of three *CCS*, a single *starting Population* in the *Grassland Cell* closest to the geometric center of either one of the 107 *Climate Cells* and one out of 18 *mowing schedules* (Table S1.4). Ignoring replicates, this results in a total of 5,778 distinct simulations runs. In the *Grassland Cell* of the starting *Population* the base *mowing schedule* named M20+00+44 always applies. Here, the first number of the schedule's name stands for early mowing calendar week 20 (day 133) and the last number for late mowing week 44 (day 301). The middle number defines the (additional) mowing weeks 22-38 of more intensive grassland management schedules (acronyms: M22-M38). All other cells receive the initially defined schedule. In that way, the starting location works as a rather undisturbed habitat of low-impact grassland mowing rendering it a fixed point for analyzing the dispersal process. Early mowing (day 133) for schedules M22-25 is omitted, because cuts should be at least six weeks apart, and late mowing (day 301) for schedules M35-38, because it us unnecessary due to slowed grassland growth and being economically ineffective for farmers (Gerling et al., 2022). Per replicate run, each simulation year was resampled randomly using ±10 years, e.g. the simulation year 2053 was determined using the set $\{2043, 2044, ..2063\}$. If the set would exceed the available data (e.g. for years $> 2080$), it is reduced accordingly.

M3: The initialization is similar to the one in M2 with some exceptions: (1) a single known population is initially placed either in spatially aggregated or fragmented grasslands of the study region (cf. Table S1.3, $reg_{init}$); (2) *Grassland Cells* are either subject to the *conventional mowing schedule* $T_{conv}$ or, with probability $p_{prot}$, to the *protective schedule* $T_{prot}$ (cf. Table S1.4), where that schedule at the initial cell is always $T_{prot}$; (3) the protected grassland probability $p_{prot} \in \{0.01, 0.02 \ldots, 0.2, 1.0\}$ (cf. Table S1.3) is defined at simulation start; (4) the timing of the *mowing schedules* is defined to be coupled to the start of the vegetation period $t_{veg}$ (Section 7.7); (5) LDD remained disabled; and (6) a number of 100 replicates were run each using a distinct *random seed*.

**Table S1.3:** List of parameters used to initialize a simulation run and to evaluate simulation results. First column: parameter name used in text. Second column: parameter symbol when used in model equations. Third column: initial / determined value(s) resp. value options used for simulation runs. Fourth column: brief description of the parameter. Parameters below double line can be varied in principle, but are constant in the presented work. M2: rows below first single line contain relevant parameters for the dispersal process. M3: rows below second single line contain parameters used for evaluating results.

| Parameter Name | Symbol | Value(s)/ Unit | Description |
|---|---|---|---|
| starting date | $t_{init}$ | $1^{st}$ Jan. of 2000, 2020, 2040 or 2060 | The date of initial time steps translated to climate data index |
| mowing day | $t_{mow}$ | none or day 134, 141,..,274 | The timing of mowing per year |
| Climate change scenario | $CCS$ | FF, MOD or BAU | Representative Concentration Pathways of $CO_2$ model |
| mowing schedule | $T_{mow}$ | see Table S1.4 | A set of mowing days per year |
| initial region | $reg_{init}$ | $\in \{aggr, frag\}$ | Definition of originating habitat (region) in terms of spatial configuration of grassland surrounding it, where aggr=aggregated and frag=fragmented |
| protected grassland probability | $p_{prot}$ | $\in \{0.01, 0.02, \ldots, 0.2, 1.0\}$ | Probability of grassland to be defined as protected habitat ($T_{mow} = T_{prot}$) at simulation start (cf. Table S1.4) |
| duration | $t_{\Delta}$ | $7,280$ or $21,840$ days | Runtime in days resp. time steps |
| habitat area | $A_{hab}$ | $250 \times 250\ m^2$ | Area of a Grassland Cell |
| initial density | $dens_{init}$ | $\{0, 0.016 \vee 0.725, 0, 0, 0\}$ $ind.\ m^{-2}$ | The initial Population density per Life Stage in individuals $m^{-2}$ |
| carrying capacity | $cap_{above}$ | $25\ ind.\ m^{-2}$ | Maximum aboveground density per square meter |
| climate cell size | $size_{clim}$ | $12,000\ m$ | Width and height of square Climate Cells |
| habitat size | $size_{hab}$ | $250\ m$ | Width and height of square habitats / Grassland Cells |
| dispersal radius | $rad_{disp}$ | $1,500\ m$ | Maximum distance covered by an individual (Griffioen, 1996) |
| base dispersal rate | $rate^{ima}_{disp}$ | $0.00595\ day^{-1}$ | Daily rate of furthest dispersing imagos (Malkus, 1997) |
| dispersal preference | $pref^{near}$ | 1 | Preference of selecting a neighbor during the dispersal process. Higher values result in selection of closer neighbors. |
| sight | $sight_{disp}$ | 0.5 | Ability to find a selected neighbor during a dispersal process |
| decay rate | $dec_{disp}$ | 0.04 | Distance dependent probability of surviving dispersal |
| inhabited status | $stat_{inh}$ | $\in \{occ, est, res\}$ | The inhabited status of a grassland habitat, where occ=occupied, est=established, res=residential |
| potential range | $rng_{pot}$ | m | The distance in meters from habitat of origin to farthest habitat (in)directly connected by $rad_{disp}$ |
| realized range | $rng_{inh}$, $inh \in stat_{inh}$ | m | The distance in meters from habitat of origin to farthest occupied / established / residential habitat |
| number of changing habitats | $\Delta n_{inh}$, $inh \in stat_{inh}$ | $\in \mathbb{N}$ | Yearly number of habitats changing their inhabited status to occupied /established /residential |
| population density | $dens_{occ}$ | $ind.\ m^2$ | Population density in $individuals\ m^2$ considering all occupied habitats in the region |

**FF=full force, MOD=moderate, BAU=business as usual, CCS=climate change scenario, aggr=aggregated, above=aboveground, cap=capacity, clim=climate, dec=decay, dens=density, disp=dispersal, est=established, frag=fragmented, hab=habitat, ima=imago, inh=inhabited, init=initial, m=meters, mow=mowing, occ=occupied, pot=potential, pref=preference, prot=protective / protected, rad=radius, reg=region, res=residential, rng=range, scen=scenario, stat=status, t=time step, veg=vegetation**

**Table S1.4:** Yearly grassland mowing schedules as applied in the simulation runs. First column gives the names of the 18 mowing schedules that encode the calendar weeks of yearly mowing occurrence divided by a plus (+) symbol. An acronym of the schedule name is provided in the second column, encoding the relevant mowing week in its name. The last three columns give the actual yearly mowing days (first day of respective calendar week) per mowing schedule. Schedules that include cells containing a dash (encoded by '00' in the respective name) only have two mowing occurrences per year, all others have three. The first mowing schedule M20+00+44 represents low-impact mowing, while more intensive mowing schedules follow in the rows below the double line.
M3: Deviating timing for dynamic mowing schedules in conventionally managed ($T_{conv}$) and protected grasslands ($T_{prot}$). Mowing occurs x days after yearly start of vegetation period $t_{veg}$ (Table S1.3).

| Schedule name | Acronym | Mowing days | | | | |
|---|---|---|---|---|---|---|
| M20+00+44 | M00 | 133 | - | 301 | | |
| M00+22+44 | M22 | - | 147 | 301 | | |
| M00+23+44 | M23 | - | 154 | 301 | | |
| M00+24+44 | M24 | - | 161 | 301 | | |
| M00+25+44 | M25 | - | 168 | 301 | | |
| M20+26+44 | M26 | 133 | 175 | 301 | | |
| M20+27+44 | M27 | 133 | 182 | 301 | | |
| M20+28+44 | M28 | 133 | 189 | 301 | | |
| M20+29+44 | M29 | 133 | 196 | 301 | | |
| M20+30+44 | M30 | 133 | 203 | 301 | | |
| M20+31+44 | M31 | 133 | 210 | 301 | | |
| M20+32+44 | M32 | 133 | 217 | 301 | | |
| M20+33+44 | M33 | 133 | 224 | 301 | | |
| M20+34+44 | M34 | 133 | 231 | 301 | | |
| M20+35+00 | M35 | 133 | 238 | - | | |
| M20+36+00 | M36 | 133 | 245 | - | | |
| M20+37+00 | M37 | 133 | 252 | - | | |
| M20+38+00 | M38 | 133 | 259 | - | | |
| conventional | $T_{conv}$ | 42 | 84 | 126 | 168 | 210 |
| protective | $T_{prot}$ | 49 | - | - | - | 217 |

### 6 Input Data

M1: As input source, time series of climate data for each *Grid Cell* are used. Each climate parameter (*ts*, *pr*, *mrso*, *smt* – cf. Leins et al. (2021), Section 2.3 and Supplement S1) is read from a single NetCDF[3] data file that stores one data point per day and coordinate. Files provided for this work contain time series from 01 January 1995 to 31 December 2080 including leap years. All data points on 29 February and 31 December are removed to achieve 364 day long years (Section 2).

M2: Climate parameters of a *Grassland Cell* are determined using up to four terrestrial *Climate Cells* in direct squared neighborhood that in terms of their geometrical center are closest to the *coordinate* of the *Grassland Cell*. The parameter values are then calculated using bilinear interpolation (Section 7.6) resulting in a two-dimensionally gradual climate data resampling of higher spatial resolution (107 to 72,969 spatial data points).

### 7 Submodels

M1: Table S1.5 gives an overview of the model processes and dynamics rates. The underlying full equations are provided in Table S1.6 and the corresponding parameters in Table S1.7.

[3]Network Common Data Form (NetCDF) library documentation: www.unidata.ucar.edu/software/netcdf/

M2: Sections 7.1.7, 7.5 and 7.6 describe the dispersal process between *Populations*, the setup of a dispersal network within a neighborhood of *Grassland Cells* and the calculation of climate values for a *Grassland Cell* using bilinear interpolation of values stemming from coarse-scale parameters of adjacent *Climate Cells*.

M3: Sections 7.1.6 and 7.1.7 include descriptions of randomly selected *mowing schedules* and the calculation of discrete amounts of dispersing individuals, respectively, while Section 7.7 introduces the temperature driven calculation of vegetation start per year and *Grassland Cell*.

#### 7.1 Update Environmental Drivers

M2: The environmental drivers are updated every time step for each *Grassland Cell*. This means here that the climate values are recalculated by bilinear interpolation (Section 7.6) and it is checked whether a *mowing schedule* takes effect.

M3: The timing of a mowing event (Section 7.1.6) can be associated with the start of the local vegetation period (Section 7.7), if defined accordingly at simulation start. Timing can thus change dynamically from year to year while slightly differing between *Grassland Cells*.

M1: Both *flow rates* and a *Cohort*'s *development progress* can change depending on the current value of environmental drivers or static conditions. Our model includes functions or equations – called *Influences* – that may be applied to achieve dynamic changes. Generally speaking, each *Influence* provides a factor that can mediate the effect of environmental conditions on the variables *dynamic flow rate* and *development progress* of a *Flow* or *Cohort*, respectively. This factor may be restricted to a minimum and maximum value ($f_{min} = 0$ and $f_{max} = 1$, if not specified otherwise) regardless of those conditions and contributes either in a multiplicative or additive way to the update of a *rate* or *progress*. The contribution to an *Influence's* current factor $f_{inf}$ by a multiplicative factor $f_{mult}$ is – as the name suggests – simply $f_{inf} \times f_{mult}$; while an additive factor $f_{add}$ contributes via the equation

$$f_{inf} + (1 - f_{inf}) \times f_{add} \tag{S1.1}$$

A factor is calculated during the update of the associated *Flow* / *Cohort*. The specific *Influences* are described below and their equations summarized in Table S1.6. *Influences* may be combined for the use in a process. The empirical basis for these equations is given below.

M1: The *Capacity Influence* is used in a logistic function that increases its factor with increasing *population density*. It restricts *Life Stages* to a maximum size by raising the mortality for large populations (Eqn. 2.2).

A *Linear Climate Influence* provides a factor that is linearly correlated with a specified climate value either in a positive or negative manner (Eqn. 2.3).

Similar to the linear correlation, an *Exponential Climate Influence* can be used to provide a factor that is exponentially correlated to a specified climate value (Eqn. 2.4).

The *Sigmoid Climate Influence* is the third option to provide a factor by correlating with a climate value (Eqn. 2.5). It allows bounding a factor to an upper and lower limit without clipping it.

**Table S1.5:** Overview of the model processes and their daily rates and equations. Processes (second column) are distinguished by Life Stage (first column) and referenced by their symbol (third column) as used in the model equations. The fourth column defines the equation and environmental drivers (Influences, f-symbols, cf. Table S1.6) used for calculating a cohort-specific dynamic rate. Equation segments marked with *s are simplifications of the iterative process of updating the flow rate described in Algorithm S1.4. Superscript letters reference the sources used to parameterize the processes and equations for the LMG (coefficients in Table S1.7): [a]Ingrisch (1983), [b]B. Schulz (pers. comm.), [c]Wingerden et al. (1991), [d]Ingrisch & Köhler (1998), [e]Helfert & Sänger (1975), [f]Helfert (1980), [g]Kriegbaum (1988), [h]Waloff (1950) , [i]Griffioen (1996), [h]Malkus (1997).

| Life Stage (symbol) | Process | Process symbol | Dynamic rate (daily) | Description |
|---|---|---|---|---|
| pre-diapause ($S^{pre}$) | mortality | $F^{pre}_{mort}$ | $dyn^{ID}_{mort} = rate^{pre}_{mort} + (1 - rate^{pre}_{mort})^{*} \times (f^{A}_{sig} \times f^{B}_{thd} + f^{C}_{mow})$ | The base mortality rate a increases in two cases: (1) humidity-driven $(f^{A}_{sig})^{a}$ if contact water $(f^{B}_{thd})^{a}$ is missing; (2) if mowing is scheduled $(f^{C}_{mow})^{b}$ |
| | development | $prog^{pre}_{ID}$ | $prog^{pre}_{ID} = \begin{cases} prog^{pre}_{ID} + f^{D}_{thd}, & if\ f^{D}_{thd} > 0 \\ 0, & \text{otherwise} \end{cases}$ | Needs three days below 10 °C in a row $(f^{D}_{thd})^{a}$ to fully develop |
| | transfer | $F^{pre}_{trans}$ | $dyn^{ID}_{trans} = \begin{cases} 1, & if\ prog^{pre}_{ID} = 1 \\ 0, & \text{otherwise} \end{cases}$ | Immediately transfers to diapause stage if fully developed |
| diapause ($S^{dia}$) | mortality | $F^{dia}_{mort}$ | $dyn^{ID}_{mort} = rate^{dia}_{mort} + (1 - rate^{dia}_{mort}) \times f^{E}_{mow}$ | The base mortality rate increases if mowing is scheduled $(f^{E}_{mow})^{b}$ |
| | development | $prog^{dia}_{ID}$ | $prog^{dia}_{ID} = \begin{cases} prog^{dia}_{ID} + \frac{f^{F}_{thd}}{2}, & if\ prog^{dia}_{ID} < 0.5, \\ prog^{dia}_{ID} + \frac{f^{G}_{thd}}{2}, & if\ prog^{dia}_{ID} \geq 0.5 \wedge f^{G}_{thd} \neq 0, \\ 0.5, & \text{otherwise} \end{cases}$ | Needs 61 days below 5 °C to break diapause $(f^{F}_{thd})^{a,c}$ and afterwards three days above 10 °C in a row $(f^{G}_{thd})^{a}$ to fully develop. Temperatures > 5 °C before diapause is broken reverse development |
| | transfer | $F^{pre}_{trans}$ | $dyn^{ID}_{trans} = \begin{cases} 1, & if\ prog^{dia}_{ID} = 1 \\ 0, & \text{otherwise} \end{cases}$ | Immediately transfers to embryo stage if fully developed |
| embryo ($S^{emb}$) | mortality | $F^{emb}_{mort}$ | $dyn^{ID}_{mort} = rate^{emb}_{mort} + (1 - rate^{emb}_{mort})^{*} \times (f^{H}_{exp} + f^{I}_{sig} \times f^{J}_{thd} + f^{K}_{mow})$ | The base mortality is defined by the temperature $(f^{H}_{exp})^{c}$. It increases in two cases: (1) humidity-driven $(f^{I}_{sig})^{a}$ if contact water $(f^{J}_{thd})^{a}$ is missing and / or (2) if mowing is scheduled $(f^{K}_{mow})^{b}$ |
| | transfer | $F^{emb}_{trans}$ | $dyn^{ID}_{trans} = rate^{emb}_{trans} \times f^{L}_{bin}(f^{M}_{exp}, f^{N}_{exp}, dens^{ID}, prog^{name}_{ID})$ | The base transfer rate is multiplied by a temperature- $(f^{M}_{exp}, f^{N}_{exp})^{c}$, progress- and the density-driven hatching probability stemming from a binomial distribution $(f^{L}_{bin})$ |
| larva ($S^{lar}$) | mortality | $F^{lar}_{mort}$ | $dyn^{ID}_{mort} = rate^{lar}_{mort} + (1 - rate^{lar}_{mort})^{*} \times (f^{O}_{cap} + f^{P}_{lin} + f^{Q}_{mow})$ | The base mortality $rate^{d}$ increases with density $(f^{O}_{cap})^{b}$ and in two additional cases: (1) the temperature $(f^{P}_{lin})$ is below 10 °C and / or (2) if mowing is scheduled $(f^{Q}_{mow})^{b}$ |
| | transfer | $F^{lar}_{trans}$ | $dyn^{ID}_{trans} = rate^{lar}_{trans} \times f^{R}_{bin}(f^{S}_{lin}, f^{T}_{lin}, dens^{ID}, prog^{lar}_{ID})$ | The base transfer rate is multiplied by a temperature- $(f^{S}_{lin}, f^{T}_{lin})^{e,f}$, progress- and density-driven maturation probability stemming from a binomial distribution $(f^{R}_{bin})$ |
| imago ($S^{ima}$) | mortality | $F^{ima}_{mort}$ | $dyn^{ID}_{mort} = rate^{ima}_{mort} + (1 - rate^{ima}_{mort})^{*} \times (f^{U}_{cap} + f^{V}_{lin} + f^{W}_{mow} + f^{X}_{mow})$ | The base mortality $rate^{g}$ increases with density $(f^{U}_{cap})^{b}$ and in three additional cases: (1) the temperature $(f^{V}_{lin})$ is below 10 °C, (2) if mowing is scheduled $(f^{W}_{mow})^{b}$ and / or (3) loss during dispersal $(f^{X}_{bif})$ |
| | reproduction | $F^{ima}_{repr}$ | $dyn^{ID}_{repr} = rate^{ima}_{repr}$ | Fixed reproduction $rate^{d,h}$ |
| | dispersal | $F^{ima}_{disp}(a,b)$ | $dyn^{ID}_{disp} = f^{Y}_{bif}$ | Dispersal rate from a to b depends on grassland cover around a, and is stochastically determined by a binomial distribution $(f^{Y}_{bif})^{i,j}$ |

**Abbreviations: bif=binominal factor, bin=binominal climate, cap=capacity, dens=density, dia=diapause, disp=dispersal, emb=embryo, exp=exponential, f=symbol of Influence function, F=Flow, ID=Cohort identifier, ima=imago, lar=larva, lin=linear, mort=mortality, mow=mowing, pre=pre-diapause, prog=progress, repr=reproduction, sig=sigmoid, t=time step, thd=threshold, trans=transfer**

**Table S1.6:** List of equations / functions applied for environmental drivers including numbering (first column), name as used in the text (second), symbol with subscript of name abbreviation (third) and brief description (last)

| Eqn# | Name | Equation / Function (symbol) | Description |
|---|---|---|---|
| (2.2) | Capacity Influence | $f_{cap} = \left(1.0 + e^{\beta^1_{cap}\left[\frac{dens^P_{above}}{A_{hab}} - \beta^2_{cap} \times cap_{above}\right]}\right)^{-1}$ | Applying carrying capacity for the above-ground population |
| (2.3) | Linear Climate Influence | $f_{lin} = \beta^1_{lin} + \beta^2_{lin} \times \omega_{clim}$ | Linear correlation with a climate parameter |
| (2.4) | Exponential Climate Influence | $f_{exp} = \beta^1_{exp} + \beta^2_{exp} \times e^{\left(\beta^3_{exp}\omega_{clim}\right)}$ | Exponential correlation with a climate parameter |
| (2.5) | Sigmoid Climate Influence | $f_{sig} = \beta^1_{sig}\left\|1 - \left(1 + e^{-\beta^2_{sig}\left[\omega_{clim} - \beta^3_{sig}\right]}\right)^{-1}\right\|$ | Sigmoid correlation with a climate parameter |
| (2.6) | Factor Threshold | $f_{thd} = \begin{cases} \beta^1_{thd}, & if\ \omega_{clim} \leq thd_{clim}, \\ \beta^2_{thd}, & \text{otherwise} \end{cases}$ | Varying factor depending on threshold exceedance |
| (2.7) | Binominal Climate | $f_{bin}\left(\mu_{\Delta t}, \sigma_{\Delta t}, dens^{ID}, prog^{name}_{ID}\right)$ | This factor is a climate-, density- and progress-driven probability stemming from a binomial distribution. See Algorithm S1.2 for details |
| (2.8) | Land Use Influence | $f_{mow} = \begin{cases} mort_{mow}, & if\ t_{mow} = t \\ 0, & \text{otherwise} \end{cases}$ | Increased mortality if mowing occurs |
| (2.9) | Binominal Factor | $f_{bif}\left(\beta^1_{bif}, dens^{ID}\right)$ | Density-driven probability stemming from a binomial distribution of a predefined mean (factor). See Algorithm S1.3 for details |

**Abbreviations: above=aboveground, bif=binominal factor, bin=binominal climate, cap=capacity, clim=climate, dens=density, exp=exponential, hab=habitat, ID=Cohort identifier, lin=linear, mort=mortality, mow=mowing, prog=progress, sig=sigmoid, t=time step, thd=treshold**

A *Factor Threshold* is used with either of the above *Influences*. It allows applying two different values depending on whether a defined climate threshold is exceeded (Eqn. 2.6). It can be used, for instance, to either enable another *Influence* (threshold exceeded, factor = 1) or disable it (threshold not exceeded, factor = 0). An iterative use of this *Influence* is possible to apply thresholds for multiple climate values.

The *Binomial Climate (M2: renamed from Factor)* is an *Influence* that depends on climate conditions and a *Cohort*'s *density* and *development progress*. It stochastically determines its factor by drawing from a binomial distribution (Eqn. 2.7, Algorithm S1.2). In other words, it defines which share of a *Cohort* population (*density*) is affected by an event (e.g. death). The binomial distribution is approximated by the cumulative distribution of a standard normal distribution. Their mean and standard deviation are calculated using one of the above *Climate Influences*. In that way, population statistics stemming from climate-driven stochastic processes can be translated to a dynamic factor like *mortality rate*.

M1: *Land Use Influence* is a timed event that increases the mortality of a *Cohort* differently depending on its *Life Stage*. In our model, land use is defined as a mowing event that occurs once a year (Eqn. 2.8). M2: The model extension adds the option to provide a *mowing schedule* that defines a set of days per year on which land use occurs.

M2: The *Binomial Factor* works similarly to the *Binomial Climate Influence* described above but deviates in two aspects. First, mean and standard deviation are calculated using a predefined probability factor only. Second, it just uses a *Cohort*'s *density* for the calculation ignoring the *development progress* (Eqn. 2.9, Algorithm S1.3).

**Algorithm S1.2** 'Binominal Climate' (M2: renamed from Factor) function to calculate a flow rate using a stochastically determined flow amount taken from a binominal distribution that is emulated by a normal distribution. The flow probability used to calculate the flow amount is taken from cumulative normal standard deviations using climate-dependent mean and standard deviation of flow duration. The resulting absolute flow rate is the ratio from flow amount to Cohort density. Finally, the applied relative flow rate takes development progress – or rather previously determined flow amount – into account to ensure flow of full density after maximum flow duration / full progress is reached.

**function** INFLUENCE('Binominal Climate', $\mu_{\Delta t}$, $\sigma_{\Delta t}$, $Cohort^{ID}$) ▷ $\mu_{\Delta t}$ and $\mu_{\Delta t}$: climate-dependent ▷ mean flow duration and standard deviation

- $density \leftarrow dens^{ID}$ ▷ get the Cohort's density
- $progress \leftarrow prog_{ID}$ ▷ get the Cohort's (development) progress
- $max_{\Delta t} \leftarrow \mu_{\Delta t} + 3 \times \sigma_{\Delta t}$ ▷ climate-dependent maximum flow duration
- $age \leftarrow max_{\Delta t} \times progress$ ▷ climate-dependent relative Cohort age
- $diff_{tod} \leftarrow frac(age - \mu_{\Delta t})\sigma_{\Delta t}$ ▷ today's age difference to mean duration
- $diff_{tom} \leftarrow frac(age + 1 - \mu_{\Delta t})\sigma_{\Delta t}$ ▷ tomorrow's age difference to mean duration
- $p_{trans} \leftarrow CDF(diff_{tom}) - CDF(diff_{tod})$ ▷ flow probability; CDF: cumulative standard normal distribution
- $\mu_{amount} \leftarrow density \times p_{trans}$ ▷ mean flow amount
- $\sigma_{amount} \leftarrow \sqrt{\mu_{amount} \times (1 - p_{trans})}$ ▷ standard deviation of flow amount
- $n_{amount} \leftarrow NORM(\mu_{amount}, \sigma_{amount})$ ▷ draw from normal distribution (NORM) to determine amount
- $r_{abs} \leftarrow fracn_{amount}density$ ▷ absolute flow rate
- $r_{rel} \leftarrow 0$ ▷ relative flow rate

```
if progress < 1 then
    r_rel ← r_abs / (1−progress)
end if
if r_rel > 1 then
    r_rel ← 1
end if
progress ← progress + r_rel
return r_rel
end function
```

**Algorithm S1.3** The 'Binominal Factor' function calculates a flow rate using a stochastically determined flow amount taken from a binominal distribution that is emulated by a normal distribution. The predefined probability factor is used directly as mean value of the distribution and indirectly to calculate the standard deviation. The resulting flow rate is the ratio of flow amount compared to Cohort density.

**function** INFLUENCE('Binominal Factor', $p_{flow}$, $Cohort^{ID}$) ▷ $p_{flow}$: predefined probability factor

- $density \leftarrow dens^{ID}$ ▷ get the Cohort's density
- $\mu_{amount} \leftarrow density \times p_{flow}$ ▷ mean flow amount
- $\sigma_{amount} \leftarrow \sqrt{\mu_{amount} \times (1 - p_{flow})}$ ▷ standard deviation of flow amount
- $n_{amount} \leftarrow NORM(\mu_{amount}, \sigma_{amount})$ ▷ draw from normal distribution (NORM) to determine amount
- $r_{flow} \leftarrow \frac{n_{amount}}{density}$ ▷ flow rate

```
if r_flow > 1 then
    r_rel ← 1
end if
return r_flow
end function
```

M1: In the following, we describe the life cycle processes of the LMG separately for each life stage and define the environmental drivers influencing them. The parametrization for the belowground life stages is mainly adapted from *temperature* experiments conducted by Wingerden et al. (1991) and *soil moisture* and *contact water* experiments by Ingrisch (1983).

For the aboveground population, data from different sources was used (Sections 7.1.4 and 7.1.5) – not all of them including LMG explicitly. The parameter values for the impact of grassland were established following personal communication with B. Schulz (Section 7.1.6).

M2: Information incorporated to estimate and validate the dispersal process (Section 7.1.7) of the LMG are an empirical 'mark and recapture' study by Malkus (1997), field studies by Griffioen (1996) and Marzelli (1994), as well as two experiments using genetic markers to measure distances between populations (Keller, 2012; Van Strien, 2013).

**Table S1.7:** List of coefficients (third column) used to parameterize the Influences' equations (second column, Table S1.6) as used to modify the processes (first column, Table S1.5) of the target species. Superscript letters reference the sources used to parameterize the processes and influences for the LMG: [a]Ingrisch (1983), [b]B. Schulz (pers. comm.), [c]Wingerden et al. (1991), [d]Ingrisch & Köhler (1998), [e]Helfert & Sänger (1975), [f]Helfert (1980), [g]Kriegbaum (1988), [h]Waloff (1950) , [i]Griffioen (1996), [h]Malkus (1997)

| Process symbol | Influences | Base rate of process and / or coefficients of influences |
|---|---|---|
| $F_{mort}^{pre}$ | <br>$f_{sig}^{A}$<br>$f_{thd}^{B}$<br>$f_{mow}^{C}$ | $(rate_{mort}^{pre} = 8.164147 \times 10^{-4})^a$<br>$(\beta_{sig}^{A1} = 0.357,\ \beta_{sig}^{A2} = 17.183,\ \beta_{sig}^{A3} = 0.703,\ \omega_{clim}^{A} = \omega_{rhug})^a$<br>$(\beta_{thd}^{B1} = 1,\ \beta_{thd}^{B2} = 0,\ \omega_{clim}^{B} = \omega_{cw},\ thd_{clim}^{B} = 0\ kg\ m^{-2})^a$<br>$(mort_{mow}^{C} = 0.05)^b$ |
| $prog_{ID}^{pre}$ | $f_{thd}^{D}$ | $(\beta_{thd}^{D1} = \frac{1}{3},\ \beta_{thd}^{D2} = 0,\ \omega_{clim}^{D} = \omega_{ts},\ thd_{clim}^{D} = 10°C)^a$ |
| $F_{mort}^{dia}$ | $f_{mow}^{E}$ | $(rate_{mort}^{pre} = 8.164147 \times 10^{-4})^a,\ (mort_{mow}^{E} = 0.05)^b$ |
| $prog_{ID}^{dia}$ | $f_{thd}^{F}$<br>$f_{thd}^{G}$ | $(\beta_{thd}^{F1} = \frac{1}{61},\ \beta_{thd}^{F2} = \frac{-1}{182},\ \omega_{clim}^{F} = \omega_{ts},\ thd_{clim}^{F} = 5°C)^{a,c}$<br>$(\beta_{thd}^{G1} = 0,\ \beta_{thd}^{G2} = \frac{1}{3},\ \omega_{clim}^{G} = \omega_{ts},\ thd_{clim}^{G} = 10°C)^a$ |
| $F_{mort}^{emb}$ | $f_{exp}^{H}$<br>$f_{sig}^{I}$<br>$f_{thd}^{J}$<br>$f_{mow}^{K}$ | $(\beta_{exp}^{H1} = 6.949 \times 10^{-3},\ \beta_{exp}^{H2} = 5.45 \times 10^{-8},\ \beta_{exp}^{H3} = 0.4743,\ \omega_{clim}^{H} = \omega_{ts})^c$<br>$(\beta_{sig}^{I1} = 0.351,\ \beta_{sig}^{I2} = 40,\ \beta_{sig}^{I3} = 0.975,\ \omega_{clim}^{I} = \omega_{rhug})^a$<br>$(\beta_{thd}^{J1} = 1,\ \beta_{thd}^{J2} = 0,\ \omega_{clim}^{J} = \omega_{cw}, thd_{clim}^{J} = 0\ kg\ m^{-2})^c$<br>$(mort_{mow}^{K} = 0.05)^b$ |
| $F_{trans}^{emb}$ | $f_{exp}^{M}$<br>$f_{exp}^{N}$ | $(\beta_{exp}^{M1} = 7.991,\ \beta_{exp}^{M2} = 1069.98,\ \beta_{exp}^{M3} = -0.2248,\ \omega_{clim}^{M} = \omega_{ts})^a$<br>$(\beta_{exp}^{N1} = 0.9251,\ \beta_{exp}^{N2} = 126.3933,\ \beta_{exp}^{N3} = -0.1978,\ \omega_{clim}^{N} = \omega_{ts})^a$ |
| $F_{mort}^{lar}$ | <br>$f_{cap}^{O}$<br>$f_{lin}^{P}$<br>$f_{mow}^{Q}$ | $(rate_{mort}^{lar} = 0.0358)^d$<br>$(\beta_{cap}^{O1} = -1.5,\ \beta_{cap}^{O2} = 0.85)$<br>$(f_{max}^{P} = 0.95,\ \beta_{lin}^{P}1 = 1.9,\ \beta_{lin}^{P2} = -0.19,\ \omega_{clim}^{P} = \omega_{ts})$<br>$(mort_{mow}^{Q} = 0.95)^b$ |
| $F_{trans}^{lar}$ | $f_{lin}^{S}$<br>$f_{lin}^{T}$ | $(\beta_{lin}^{S1} = 83,\ \beta_{lin}^{S2} = -1.679,\ \omega_{clim}^{S} = \omega_{ts})^{e,f}$<br>$(\beta_{lin}^{T1} = -2.2188,\ \beta_{exp}^{T2} = 0.2188,\ \omega_{clim}^{T} = \omega_{ts})^{e,f}$ |
| $F_{mort}^{ima}$ | <br>$f_{cap}^{U}$<br>$f_{lin}^{V}$<br>$f_{mow}^{W}$<br>$f_{bif}^{X}$ | $(rate_{mort}^{ima} = 0.0475)^g$<br>$(\beta_{cap}^{U1} = -1.5,\ \beta_{cap}^{U2} = 0.85)$<br>$(f_{max}^{V} = 0.95,\ \beta_{lin}^{V1} = 1.9,\ \beta_{lin}^{V2} = -0.19,\ \omega_{clim}^{V} = \omega_{ts})$<br>$(mort_{mow}^{W} = 0.95)^b$<br>$(\beta_{bif}^{X1} = mort_{disp}^{ima}$, see Eqn. S1.16) |
| $F_{repr}^{ima}$ | | $(rate_{repr}^{ima} = 1.3)^{d,h}$ |
| $F_{disp}^{ima}$ | $f_{bif}^{Y}$ | $(\beta_{bif}^{Y1} = rate_{disp}^{ima}(a,b)$, see Eqn. S1.11$)^{i,j}$ |

**Abbreviations: bif=binominal factor, dia=diapause, disp=dispersal, emb=embryo, exp=exponential, f=symbol of Influence function, F=Flow, ID=Cohort identifier, ima=imago, lar=larva, lin=linear, mort=mortality, mow=mowing, pre=pre-diapause, prog=progress, repr=reproduction, sig=sigmoid, thd=treshold, trans=transfer**

M1: The values for *base rates* and the *coefficients* used to parameterize the *Influences* as applied for the processes of the target species are listed in Table S1.7. A verbal description on their usage for the target species is given in the following subsections.

##### 7.1.1 Pre-diapause Life Stage

M1: The first sub-stage of the belowground population represents the LMG's clutch after oviposition and before diapause. Following the study of Ingrisch (1983), we define that it requires three consecutive days below a *temperature* of 10 °C to start diapause thus develop into the next life stage (*Factor Threshold*, Eqn. 2.6). The intrinsic *mortality rate* rapidly increases if the eggs experience pre-winter drought stress caused by missing *contact water* (Ingrisch, 1983) (*Factor Threshold*, Eqn. 2.6). The increased mortality is calculated using the *Sigmoid Climate Influence* (Eqn. 2.5) driven by *humidity*.

##### 7.1.2 Diapause Life Stage

M1: The diapause life stage occurs mainly during winter and is mostly unaffected by climate conditions in terms of our model. Therefore, the intrinsic *mortality rate* remains constant. We defined that it needs to experience a total of 61 days below 5 °C to break diapause (*Factor Threshold*, Eqn. 2.6). By that, we followed the cold treatments in the studies by Ingrisch (1983) and Wingerden et al. (1991), though with a shorter period of time to avoid stagnation in slightly warmer winters. Additionally, a too early indication of spring is prevented by defining that the life stage requires three consecutive days of at least 10 °C to develop into the next life stage (*Factor Threshold*, Eqn. 2.6). The diapause life stage can have multiple *Cohorts*, i.e., one for each consecutive year of unsuitable conditions (diapause not broken).

##### 7.1.3 Embryo Life Stage

M1: This life stage is the most complex in our model. Mortality is influenced by three climate parameters while the temperature-driven hatching process (transfer to larval stage) is stochastically determined by the stage's *density* and *development progress*. Following Wingerden et al. (1991) we established the *mortality rate* using the fact that a *temperature* of 22.2 °C yields the highest hatching success rate (∼ 82 %) while the rate drops with both lower and higher *temperatures* (*Exponential Climate Influence*, Eqn. 2.4). At the same time higher *temperatures* decrease the mean hatching time from 45 days at 15.0 °C to 8 days at 37.5 °C (*Exponential Climate Influence*, Eqn. 2.4). In our model, the *base mortality rate* is in fact mainly determined by *temperature*. Similar to the pre-diapause life stage it can further increase in the case of post-winter drought stress caused by missing *contact water* (Ingrisch, 1983) (*Factor Threshold*, Eqn. 2.6). This additional mortality is calculated using the *Sigmoid Climate Influence* (Eqn. 2.5) driven by *humidity*. As mentioned above, the timing and amount of hatching eggs is stochastically determined (*Binomial Climate* [M2: renamed from Factor], Eqn. 2.7). The hatching probability increases with *development progress* and higher *temperatures*. Mean and standard deviation defining the binomial distribution to draw the probability from are factors that are correlated with the *temperature* (*Exponential Climate Influence*, Eqn. 2.4).

##### 7.1.4 Larva Life Stage

M1: The larval life stage has multiple *Cohorts*, i.e., one for each hatching day per year of the previous embryo stage. These larva *Cohorts* are processed independently. High *temperatures* increase the development speed of larvae. We used the development traits found in three related grasshopper species (Helfert & Sänger, 1975; Helfert, 1980) to calculate temperature-driven mean and standard deviation (*Linear Climate Influence*, Eqn. 2.3). Both are then applied to stochastically determine the density- and progress-dependent transfer (or rather development) of a larva *Cohort* to the imago life stage (*Binomial Climate* [M2: renamed from Factor], Eqn. 2.7). The larva *base mortality rate* was adopted by averaging the parameters of two related grasshopper species (Ingrisch & Köhler, 1998, Tab. 16). It increases with *aboveground population density* (*Capacity Influence*, Eqn. 2.2) allowing a maximum of 25 individuals $m^{-2}$ (B. Schulz, pers. comm.) and by definition with *temperatures* below 10 °C (*Linear Climate Influence*, Eqn. 2.3).

##### 7.1.5 Imago Life Stage

M1: Similar to the larval stage, the *base mortality rate* for the imago life stage was adopted using the daily survival rates of four related grasshopper species determined by Kriegbaum (1988). It increases with *aboveground population density* (*Capacity Influence*, Eqn. 2.2) allowing a maximum of 25 individuals $m^{-2}$ (B. Schulz, pers. comm.) and by definition with *temperatures* below 10 °C (*Linear Climate Influence*, Eqn. 2.3). The daily *transfer* or rather *oviposition rate* of 1.3 was defined including several considerations. First of all, an LMG egg pod contains 11-14 eggs (Waloff, 1950) and reproduction is delayed up to two weeks after maturation (Ingrisch & Köhler, 1998). Furthermore, we assumed that half of the LMG population is female and that a female lays up to three egg pods during a lifetime of 60 days. In terms of our model, oviposition is not influenced by any external drivers.

##### 7.1.6 Land Use

M1: As mentioned above, land use in our model is a representation of grassland mowing that occurs once per year on the first day of the same calendar week. It affects the belowground population (life stages 1-3) less severely than the aboveground population (4-5) by increasing the stage's *mortality rate* additively by 0.05 (below ground) and 0.95 (above ground), respectively (B. Schulz, pers. comm.).

M2: Disturbance through land use occurs on 2-3 days per simulation year depending on initially defined *mowing schedule* (Table S1.4).

M3: Depending on the protected grassland probability $p_{prot}$, the *mowing schedule* occurring in each *Grassland Cell* is determined randomly at simulation start. That is, either the conventional schedule $T_{conv}$ with five cuts or the protective schedule $T_{conv}$ with two cuts per year (cf. Table S1.4). Depending on the schedules' definitions the cuts occur $x$ days after the start of a year's vegetation period (Section 7.7).

##### 7.1.7 Dispersal

M2: In terms of the model extension, the dispersal rate from the imago *Life Stage* of *Population* $P_a$ to the same stage of each neighboring *Population* $P_n \in N_a$ is stochastically determined (*Binomial Factor*, Eqn. 2.9) every time step using a *base dispersal rate* ($rate_{disp}^{ima} = 0.00595\ day^{-1}$) and a pre-calculated *dispersal probability* (Eqn. S1.11). We determined the base dispersal rate by defining those individuals as dispersers that traveled the largest distance during one day (1 out of 168) in a 'mark and recapture' study by Malkus (1997). The *dispersal mortality rate* is calculated using the inverse of the dispersal rate (Eqn. S1.16). Following the maximum covered distance of LMG individuals described by Griffioen (1996), dispersal is defined to remain within a radius ($rad_{disp}$) of $1,500\ m$, in principle. In regions of low grassland cover, however, dispersal outside this radius (LDD) can occur to account for the LMG's flight ability (Sörens, 1996). Section 7.5 describes in detail, how the neighborhood (both inside and outside of the dispersal radius) of each *Population* is defined and the values of the dispersal process are determined.

M3: To prevent unrealistic low population size in newly inhabited cells, the amount of dispersing individuals is truncated to discrete numbers in terms of habitat size $A_{hab}$. Discrete here means that the size of one individual is defined as $\frac{1}{A_{hab}}\ ind.\ m^{-2}$ and the calculated dispersing amount must be a manifold of this individual size. Dispersing amount below the next largest manifold remains in the *life stage of origin*.

#### 7.2 Flow Update

M1: During 'Flow update', the state variables *total flow amount*, *current flow amount* and *dynamic flow rate* (Table S1.5) are iteratively recalculated using the *Cohorts* associated with the *life stage of origin* (Algorithm S1.4). If there are any external *Influences* (Section 7.1) associated with this *Flow*, they change the *dynamic flow rate* depending on their *type*.

#### 7.3 Life Stage Update

M1: The submodel 'Life stage update' (Algorithm S1.5) calculates a *Life Stage*'s *total density* from the *density* of its subordinate *Cohorts* and its *gain* from the incoming *transfer Flow* of its preceding *Life Stage*. The *gain* is then used to determine whether to create a new *Cohort* from it or add it to an existing *Cohort*. Furthermore, overaged *Cohorts* or such of *density* below the minimum are erased.

**Algorithm S1.4** Submodel 'Flow update'. All associated Flows of the defined Population are iteratively updated by this submodel. Sequence of update is irrelevant. Algorithm describes the update of a single Flow. Equations of influence functions are listed in Table S1-5.

```
function SUBMODEL('Flow update')
    amount ← 0                                                        ▷ total flow amount
    origin ← Flow_origin                                              ▷ Flow's Life Stage of origin
    for all Cohort^ID ∈ origin_Cohorts do
        amount^ID ← 0                                                 ▷ Cohort's flow amount
        rate^ID ← 0                                                   ▷ Cohort's flow rate
        progress ← prog_ID                                            ▷ Cohort's (development) progress
        if Flow_type = 'mortality' OR progress ≥ 1 then
            if Flow_type = 'mortality' AND age^ID ≥ age^max_origin then
                amount^ID ← 1
            else
                rate^ID ← Flow_rate                                   ▷ set to Flow's base rate
                for all Influence ∈ Flow_Influences do
                    factor_Influence ← SUBMODEL('Influence')
                    if Influence_type = 'additive' then
                        rate^ID ← rate^ID + factor_Influence × (1 − rate^ID)
                    else if Influence_type = 'multiplicative' then
                        rate^ID ← rate^ID × factor_Influence
                    end if
                end for
            end if
        end if
    end for
    amount^ID ← rate^ID × density^ID
    amount ← amount + amount^ID
end function
```

**Algorithm S1.5** Submodel 'Life Stage update'. All associated Life Stages of a Population are iteratively updated using this submodel. The pseudocode describes the update of a single Life Stage.

```
function SUBMODEL('Life Stage update')
    density ← 0                                                       ▷ total density
    gain ← 0                                                          ▷ total density gain
    created_Cohort ← false                                            ▷ was a new Cohort created?
    for all Flow ∈ Stage_input do                                     ▷ loop over incoming Flows of (Life) Stage
        gain ← gain + Flow_amount                                     ▷ add total flow amount to gain
        if gain ≥ density^min_Stage then
            if Stage_Cohorts = ∅ OR multi^Cohort_Stage = true then    ▷ empty set of Cohorts or multiple allowed
                Cohort^gaining ← CREATE COHORT(gain)                  ▷ Cohort with gain as initial density
                created_Cohort ← true
            else
                Cohort^gaining ← Stage_Cohorts(1)                     ▷ get Cohort from top of Stage's Cohorts
                density^gaining ← density^gaining + gain
            end if
        end if
    end for
    for all Cohort^ID ∈ Stage_Cohorts do
        RUN SUBMODEL('Cohort update')
        if density^ID < density^min_Stage then
            remove Cohort^ID from Stage_Cohorts
        else
            density ← density + density^ID
        end if
    end for
    if created_Cohort = true then
        Stage_Cohorts(1) ← Cohort^gaining                             ▷ put new Cohort on top of Stage's Cohorts
    end if
end function
```

#### 7.4 Cohort Update

M1: During 'Cohort update', *density* and *development progress* of a *Cohort* a recalculated (Algorithm S1.6). First, *loss* of *density* is calculated using the outgoing *transfer Flow* of the *Cohort's* parent *Life Stage*. Then, *gain* (if any) is added to the *density* and reset to zero afterwards. Finally, the *development progress* is increased using external influences, if there were defined any. Otherwise it is set to a value of 1.

**Algorithm S1.6** Submodel 'Cohort update'. All associated Cohorts of the defined Life Stage are iteratively updated by this submodel. Sequence of update is irrelevant. Pseudocode describes the update of a single Cohort. Equations of influence functions are listed in Table S1-5.

```
function SUBMODEL('Cohort update', ID)                          ▷ using ID of updating Cohort
    loss ← 0                                                     ▷ total density loss
    parent ← Cohort_parent                                       ▷ Cohort's parent Life Stage
    for all Flow ∈ parent_output do                              ▷ loop over outgoing Flows of parent Life Stage
        if Flow_type ≠ 'reproduction' then
            loss ← loss + Flow^ID_amount                         ▷ add Cohort-specific flow amount to loss
        end if
    end for
    density ← density + (gain − loss)                            ▷ update Cohort density with potential gain and loss
    gain ← 0                                                     ▷ reset (externally updated) gain
    progress_temp ← 1                                            ▷ temporary buffer for potential development progress
    for all Influence ∈ Development_Influences do                ▷ loop over development Influences, if any
        factor_Influence ← SUBMODEL('Influence')
        progress_temp ← factor_Influence × progress_temp
    end for
    progress ← progress + progress_temp
    if progress < 0 then
        progress ← 0
    else if progress > 1 then
        progress ← 1
    end if
end function
```

#### 7.5 Dispersal Setup

M2: The submodel 'Dispersal setup' is called every time the *life stage of destination* of an empty *Population* is subject in a non-zero *dispersal Flow* (in terms of *flow amount*), in other words, if dispersal is directed to an *uninhabited Grassland Cell* (for the first time). In this case, the submodel locates all *Populations* $P_b \in N_a \subset R$ in the study region R belonging to the sub region or neighborhood $N_a$ of *Population* $P_a$ in the now *inhabited* cell $G_a$ and establishes a connection to each of them by creating a *dispersal Flow* between the imago *Life Stages* of the formerly empty *Population* (*life stage of origin*) and the respective neighboring *Population* (*life stage of destination*). *Populations* considered a neighbor $P_b \in N_a$ are determined in two ways: (1) all *Populations* inside a predefined dispersal radius (home range); and (2) the nearest *Population* in either one of eight cardinal directions [North (N), Northeast (NE), East (E), Southeast (SE), South (S), Southwest (SW), West (W), Northwest (NW)], in case there are no cells found inside the home range in this direction (LDD). Identifying *Populations* inside the home range is straight forward using the Euclidean distance $dist_{a,b}$ [in meters] between two *Grassland Cells* $G_a$ and $G_b$:

$$dist_{a,b} = \sqrt{(coord_x^a - coord_x^b)^2 + (coord_y^a - coord_y^b)^2} \times size_{hab} \tag{S1.10}$$

If $dist_{a,b} \leq rad_{disp}$ (Table S1.3), the cells belong to the same neighborhood. Finding potential neighbors for LDD into one of the cardinal directions $DIR = N, NE, E, SE, S, SW, W, NW$ requires additional information about the surroundings of the source *Grassland Cell* $G_a$. If none of the *Populations* $P_c \in N_a^{dir}$ fulfills the home range constraint $dist_{a,c} \leq rad_{disp}$, the nearest cell $P_{LDD} \in N_{a,LDD}^{dir} \subset N_a^{dir}$ is selected as long distance neighbor in direction $dir$. Here, $N_a^{dir}$ represents a set of all *Populations* located within a 90-degree angle in direction $dir \in DIR$ of cell $G_a$ and $N_{a,LDD}^{dir}$ a subset of slightly narrowed angle outside the dispersal radius. Using this method, potential long distance neighbors are identified for all directions of $DIR$. Figure 5 in Leins et al. (2022) geometrically illustrates the selection of LDD neighbors. A detailed definition of cells belonging to each of the narrowed subsets will follow below. Having identified all *Populations* belonging to the neighborhood $N_a$, the *flow rate* $rate_{disp}^{ima}(a,b)$ for each dispersal *Flow* originating in $G_a$ and destined in $G_b$ is determined using the base dispersal rate defined for the *Life Stage* of interest (here, $rate_{disp}^{ima}$ for the LMG's imago stage) and a species-specific *dispersal probability* $p_{a,b}^{disp}$ (Eqn. S1.11):

$$rate_{disp}^{ima}(a,b) = rate_{disp}^{ima} \times p_{a,b}^{disp} \tag{S1.11}$$

$$p_{a,b}^{disp} = pref_{a,b} \times p_{a,b}^{find} \times p_{a,b}^{surv} \tag{S1.12}$$

The *dispersal probability* itself (Eqn. S1.12) is calculated using a *preference factor* to select the destined *Population* depending on the distance to all neighbors (Eqn. S1.13), a probability to find the selected neighbor during the dispersal process (Eqn. S1.14) and a probability to survive the dispersal (Eqn. S1.15):

$$pref_{a,b} = \frac{1/dist_{a,b}^{pref^{near}}}{\sum_{P_n \in N_a} dist_{a,n}^{pref^{near}}} \tag{S1.13}$$

$$p_{a,b}^{find} = \left(rGrass_{a,b}^{at}\right)^{1-sight_{disp}} \tag{S1.14}$$

$$p_{a,b}^{surv} = \exp^{-decay_{disp} \times (1 - rGrass_{a,b}^{to}) \times dist_{a,b}} \tag{S1.15}$$

Table S1.3 contains the parameter values used to define the dispersal process of the LMG and their usage is described in the following. The size of the parameter $pref^{near} \geq 0$ in Eqn. S1.13 defines to what extent nearby cells are preferred over more distant ones, where a value of zero leads to equal preference independent of the distance. Finding a selected neighbor (Eqn. S1.14) depends on the relative grassland cover $rGrass_{a,b}^{at}$ in the same distance as the destined cell (i.e., the actual number of *Grassland Cells* relative to the potential number), and the ability of the species to locate suitable neighbors given by the parameter $sight_{disp} \in [0,1]$, where a value of zero defines a success rate equal to the grassland ratio and a value of one a 100 % success rate. The probability to survive the dispersal process (Eqn. S1.15) depends on the relative grassland cover $rGrass_{a,b}^{to}$ in the perimeter of a radius lower than the distance to the destined cell and a distance dependent decay rate $decay_{disp} \in [0,1]$, where a value of zero equals 100 % and a value of one equals minimum survival probability. Furthermore, the dispersers are subject to dispersal mortality which is the difference of the sum of all the dispersal probabilities multiplied by the *base dispersal rate*:

$$mort_{disp}^{ima} = \left(1 - \sum_{P_n \in N_a} p_{a,n}^{disp}\right) \times rate_{disp}^{ima} \tag{S1.16}$$

Finally, the eight sub regions $N_{a,LDD}^{dir} \subset R$ representing all cells found outside the dispersal radius in the direction $dir \in DIR$ of an originating *Grassland Cell* $G_a$ are determined as follows:

$$N_a^N = \{\forall G_n \in R \backslash \{G_a\} \mid dist_{a,n}^x < dist_{a,n}^y \wedge coord_y^n < coord_y^a \wedge dist_{a,n} > rad_{disp}\} \tag{S1.17}$$
$$N_a^{NE} = \{\forall G_n \in R \backslash \{G_a\} \mid coord_x^n > coord_x^a \wedge coord_y^n < coord_y^a \wedge dist_{a,n} > rad_{disp}\} \tag{S1.18}$$
$$N_a^E = \{\forall G_n \in R \backslash \{G_a\} \mid coord_x^n > coord_x^a \wedge dist_{a,n}^y < dist_{a,n}^x \wedge dist_{a,n} > rad_{disp}\} \tag{S1.19}$$
$$N_a^{SE} = \{\forall G_n \in R \backslash \{G_a\} \mid coord_x^n > coord_x^A \wedge coord_y^n > coord_y^a \wedge dist_{a,n} > rad_{disp}\} \tag{S1.20}$$
$$N_a^S = \{\forall G_n \in R \backslash \{G_a\} \mid dist_{a,n}^x < dist_{a,n}^y \wedge coord_y^a > coord_y^a \wedge dist_{a,n} > rad_{disp}\} \tag{S1.21}$$
$$N_a^{SW} = \{\forall G_n \in R \backslash \{G_a\} \mid coord_x^n < coord_x^a \wedge coord_y^a > coord_y^a \wedge dist_{a,n} > rad_{disp}\} \tag{S1.22}$$
$$N_a^W = \{\forall G_n \in R \backslash \{G_a\} \mid coord_x^n < coord_x^a \wedge dist_{a,n}^y < dist_{a,n}^x \wedge dist_{a,n} > rad_{disp}\} \tag{S1.23}$$
$$N_a^{NW} = \{\forall G_n \in R \backslash \{G_a\} \mid coord_x^n < coord_x^a \wedge coord_y^n < coord_y^a \wedge dist_{a,n} > rad_{disp}\} \tag{S1.24}$$

We defined the sub regions in such a way that each of them overlaps with both neighbors in clockwise and counterclockwise direction to minimize LDD to cases where the area in direction of interest is largely empty within the home range. Figure 5 in Leins et al. (2022) visualizes the areas belonging to either of the sub regions.

##### 7.6 Bilinear Climate Interpolation

M2: This submodel is called every time step to achieve heterogeneous, gradual values at the location of each *Grassland Cell* using bilinear interpolation of the climate data from the closest adjacent neighbors. This is done by weighing the distances from a *Grassland Cell* $G_a$ to the center of the (up to) four *Climate Cells* $\{\Omega_{a,NE}, \Omega_{a,SE}, \Omega_{a,SW}, \Omega_{a,NW}\}$ into secondary cardinal directions of $G_a$. The resulting bilinear weights $w_{bilin}^{a,dir}$ are multiplied with their respective climate values $\omega_{clim}^{a,dir}$ and then summed to achieve the value at $G_a$. The interpolated climate value $\omega_{clim}^a$ for cell $G_a$ is then calculated as follows.

$$\omega_{clim}^a = \sum_{dir \in DIR_{sec}} w_{bilin}^{a,dir} \times \omega_{clim}^{a,dir} \tag{S1.25}$$
$$w_{bilin}^{a,dir} = 1 - \frac{(size_{clim} - dist_{a,dir}^x) \times (size_{clim} - dist_{a,dir}^y)}{(size_{clim})^2} \tag{S1.26}$$
$$dist_{a,dir}^{xy} = |coord_{xy}^a - center_{xy}^{a,dir}| \times size_{hab} \tag{S1.27}$$

Here, $DIR_{sec} \subset DIR$ (see Section 7.5) are the secondary cardinal directions $\{NE, SE, SW, NW\}$ and $\omega_{clim}^{a,dir}$ is the projected value in the *Climate Cell* $\Omega_{a,dir}$ into direction dir of $G_a$. Parameters $size_{clim}$ of a *Climate Cell* and $size_{hab}$ of a *Grassland Cell* were introduced in Table S1.3. The value $dist_{a,dir}^{xy}$ for the distances in x- or y-direction are calculated using the respective *coordinate* of the geometric center of a *Climate Cell* ($center_{x,y}^{a,dir}$, Table S1.3). Figure 4 in Leins et al. (2022) illustrates the calculation of the weights for a single *Grassland Cell* using a simplified geometric example. Please refer to Leins et al. (2022), Supplement S4 for a description of the mapping between *Climate* and *Grassland Cells*, and a reference to the calculated weights.

#### 7.7 Start of Vegetation Period

The start of the vegetation period $t_{veg}$ in a *Climate* or *Grassland Cell* is calculated using the *surface temperature* $\omega_{ts}$. Following Gerling et al. (2020), this period starts when the *yearly temperature sum* surpasses 200.0 °C. Adapting their calculation to the *surface temperature* $\omega_{ts}$ used in the present model gives the following equation:

$$
\begin{aligned}
sum_{ts} &= \sum_{i=1}^{I} (x \times \omega_{ts}^{i}) \forall I \in \{1, 2, \ldots, 364\} \; until \; sum_{ts} \geq 200.0\,°C, \\
x &= \begin{cases} 0.5, & if \; 1 \leq i \leq 31 \\ 0.75, & if \; 32 \leq i \leq 59 \\ 1.0 & if \; i \geq 60 \end{cases}, \\
\omega_{ts}^{i} &= \begin{cases} 0, & if \; \omega_{ts}^{i} < 0 \\ \omega_{ts}^{i}, & \text{otherwise} \end{cases}
\end{aligned} \tag{S1.28}
$$

Here, $sum_{ts}$ is the *summed surface temperature*, $\omega_{ts}^{i}$ is the *mean surface temperature* (ignoring negative values) on day $i$ of a year, and $x$ is a weight including the *temperature* values of January and February with only 50 % and 75 % of their extent. The value of $t_{veg}$ equals the day $i$ where $t_{sum}$ reaches 200.0 °C.
